# Defective Trafficking of Annexins to the Site of Injury in ANO5-Knockout Muscle Fibers

**DOI:** 10.1101/2020.05.22.110825

**Authors:** Steven J. Foltz, YuanYuan Cui, Hyojung J. Choo, H. Criss Hartzell

**Affiliations:** Department of Cell Biology, Emory University School of Medicine, Atlanta, GA 30322

**Keywords:** Annexins, membrane repair, muscular dystrophy, ANO5/TMEM16E, phospholipid scrambling

## Abstract

Mutations in *ANO5* (*TMEM16E*) cause limb-girdle muscular dystrophy R12 (limb-girdle muscular dystrophy type 2L). Recent evidence implicates defective plasma membrane repair as a likely mechanism for the disorder. Here, we probe the ANO5-dependency of the membrane repair pathway using a laser wounding assay in *Ano5* knockout mouse muscle fibers. Wounded myofibers from *Ano5* knockout mice exhibit delayed membrane resealing relative to wild type fibers as revealed by an increased uptake of the membrane-impermeant FM1-43 dye and a prolonged elevation of intracellular Ca^2+^. The trafficking of several annexin proteins, which together form a cap at the site of injury, is altered in *Ano5* knockout fibers. Annexin A2 accumulates at the wound to nearly twice the level observed in WT fibers, while annexin A6 accumulation is substantially inhibited in the absence of ANO5. Furthermore, trafficking of annexins A1 and A5 to the cap is decreased in the *Ano5* knockout. These changes are correlated with an alteration in the fine structure of the annexin repair cap and the shedding of annexin-positive extracellular vesicles. Our results suggest that the meticulous coordination of the annexin repair machinery required to effectively reseal wounded sarcolemma is disrupted in *Ano5* knockout mice. ANO5 is a putative phospholipid scramblase, responsible for exposure of intracellular phospholipids to the extracellular leaflet of the plasma membrane. However, because the membrane repair defect is rescued by overexpression of wild type ANO5 or a scramblase-defective mutant, we suggest that ANO5-mediated phospholipid scrambling is not essential for membrane repair.

**Significance Statement:** Mutations in *ANO5/TMEM16E* cause myopathies of variable severity, with some patients losing ambulation entirely. Unfortunately, relatively little is known about the function of ANO5 at the protein level, but it has been suggested that ANO5 plays a role in the repair of injured muscle plasma membranes. Here, we investigate the mechanism of ANO5-mediated repair and find that annexin proteins, which in normal muscle form a cap to seal wounds, traffic abnormally to the cap when ANO5 is not expressed. Muscle fibers lacking ANO5 reseal more slowly and thus are exposed to prolonged intracellular calcium elevation that can damage the fibers. Our findings contribute to the growing literature implicating failed repair as a probable pathogenic mechanism in patients with *ANO5* mutations.

## Introduction

Recessive mutations in the *ANO5* gene are responsible for a spectrum of myopathies with variable severity. These disorders, which are characterized by muscle weakness and atrophy, include limb girdle muscular dystrophy R12 (LGMDR12/LGMD2L), Miyoshi muscular dystrophy type 3 (MMD3), and muscle weakness with myalgia and rhabdomyolysis (1–4). Since *ANO5*-myopathies were discovered in 2010 (1), >70 *ANO5* variants have been reported to be pathogenic (https://www.ncbi.nlm.nih.gov/clinvar/). In a recent screen of 35 LGMD genes in 4656 clinically-suspected LGMD patients, *ANO5* was the 4^th^ most likely contributor to LGMD phenotypes in the US population (5). Anoctamin-5 (ANO5) protein expression is absent in patients with the founder mutation (c.191dupA, p.Asn64Lysfs*15) (2, 6, 7) and ANO5 protein is reduced in patients with point mutations (e.g. c.2101A>G, p.N701D and c.2272C>T, p.R758C) (7, 8). Whereas ANO5-myopathies are inherited in a recessive manner, dominant (probably gain of function) *ANO5* mutations are associated with gnathodiaphyseal dysplasia, a skeletal syndrome characterized by cementosseous lesions of the jaw, osteomyelitis, bone fragility (9, 10) and dental tumors (11–13).

ANO5 is a member of the TMEM16/Anoctamin family that includes ion channels and phospholipid scramblases (14–16). Within this family, ANO5 is most closely related to the phospholipid scramblase (PLSase) ANO6 (10, 17–19). Phospholipid scrambling (PLS) disrupts the asymmetric distribution of phospholipids that normally exists between the leaflets of the plasma membrane (20). A consequence of PLS is the exposure of anionic phospholipids, such as phosphatidylserine (PtdSer), which are normally sequestered on the intracellular face of the plasma membrane, to the extracellular space (14, 20). We and others have observed that heterologous expression of ANO5 in HEK cells induces PLS (21–23). Furthermore, phospholipid scrambling is absent in muscle precursor cells from ANO5-KO mice despite expression of ANO6 (21), suggesting a unique, muscle-specific role for ANO5-mediated phospholipid scrambling. However, the pathologic mechanism of *ANO5* myopathies is poorly understood and it remains unclear whether ANO5-linked disorders are a consequence of defective PLS.

One suggestion is that *ANO5* myopathies are caused by defects in the ability of muscle fibers to self-repair (1, 8, 24, 25). Skeletal muscle endures considerable mechanical stress even during normal use, which can produce small tears in the plasma membrane (sarcolemma) (26). Healthy muscle has robust mechanisms to limit damage by repairing torn sarcolemma through the coordinated interplay of various proteins, lipids, organelles, and small molecules (27–31). Disrupted, slow, or incomplete resealing results in prolonged Ca^2+^ entry with consequent sustained activation of calpain endopeptidases and phospholipases that have potential to produce long-lasting muscle damage (32). Repair mechanisms are thought to be disrupted in several muscular dystrophies, most notably limb-girdle muscular dystrophy R2 (LGMDR2/LGMD2B) and Miyoshi muscular dystrophy type 1 (MMD1) (33), which are caused by mutations in the dysferlin (<u>*DYSF*</u>) gene (34, 35). We have shown that disruption of the *Ano5* gene in mice (herein ANO5-KO) causes a muscle phenotype that includes defective cell membrane repair (21, 36). Recently, the pathology of LGMDR12 has been tied to defective membrane repair through a mechanism involving ANO5-regulated Ca^2+^ handling by the endoplasmic reticulum (8). While ANO5 myopathies share features with dysferlinopathies, ANO5 and dysferlin probably have non-overlapping roles in the repair pathway because ANO5 overexpression in dysferlin-null mice does not rescue the repair defect (25).

One of the most rapid responses to membrane injury is the Ca^2+^-dependent accumulation of annexins (ANX) A1, A2, A4, A5, A6, and A11 at the wound (37–46). Moreover, repair *in vitro* and *in vivo* is enhanced by overexpression of ANXA1, A2, or A6 (41) or extracellular application of ANXA5 or A6 protein (44). These findings demonstrate that annexins play key roles in the repair process. The annexins are known to perform a spectrum of diverse membrane-related functions that range from membrane stabilization to membrane deformation and promotion of fusion (46, 47). All annexins share a basic ability to bind anionic phospholipids (phosphatidylserine and phosphatidylinositols) in a Ca^2+^-dependent manner, but their differing Ca^2+^ affinities, binding partners, and quaternary structures confer unique biological roles (46, 47). The recruitment of various annexins to the lesion occurs sequentially and in at least some cases interdependently (40, 42, 46), suggesting that they may play non-redundant parts in the repair orchestra. Unlike annexins, other major repair proteins may act Ca^2+^-independently. For example, dysferlin can bind to phospholipids through a C-terminal polybasic motif independent of its Ca^2+^-binding domains (43, 48) and the accumulation of the E3 ligase MG53 is dependent on oxidative oligomerization and Zn^2+^ (49, 50). Accumulation of these repair proteins at the lesion coincides with local enrichment of PtdSer at the plasma membrane adjacent to the lesion or in membrane “patches” that arise during repair (40, 43). This PtdSer enrichment is intriguing partly because the annexins and dysferlin both bind PtdSer, but the mechanism underlying PtdSer externalization is not known. It is possible that it could involve either membrane trafficking or phospholipid scrambling, the latter of which would provide an attractive link to ANO5 in the repair process.

In this paper, we investigate the role of ANO5 in skeletal muscle membrane repair. We show that ANO5 rapidly appears at the plasma membrane of muscle fibers damaged with a laser pulse. Furthermore, loss of ANO5 is associated with abnormal trafficking of several annexin proteins to the wound. However, contrary to our initial hypothesis, we find that PtdSer exposure during repair is not ANO5-dependent and that ANO5-mediated PLS is dispensable for repair. Lastly, we demonstrate that a patient-associated point mutation in ANO5 results in defective repair in primary human myotubes, implicating defective membrane repair in the development of LGMDR12.

## Results

### ANO5-deficient myofibers are defective in membrane repair

A standard method for studying membrane repair involves rupturing the cell membrane with an intense laser pulse (33, 51). We chose two complementary approaches to evaluate the kinetics of plasma membrane resealing after a laser pulse. First, muscle fibers from WT or ANO5-KO mice were loaded with the Ca^2+^-sensitive dye Cal-520 (K_d_ = 320nM) and the Ca^2+^ signal was measured after laser injury. Upon membrane damage, extracellular Ca^2+^ entered myofibers, resulting in a transient increase in Cal-520 fluorescence. Cal-520 fluorescence intensity in KO fibers was significantly elevated relative to WT beginning 20 s post-injury (Fig. 1A, B). In WT fibers, maximal fluorescence intensity was observed 20 s after injury and then declined with a half-time t/2 = 46 s to reach a plateau level that was ~32.8% greater than baseline 7 minutes after injury. In contrast, in ANO5-KO fibers Ca^2+^ transients peaked at 28 s post-injury and maximal Cal-520 fluorescence (normalized to initial fluorescence intensity) was significantly higher than in WT fibers. Fluorescence intensity declined with a half-time t/2 = 42 s to a final value 61% greater than baseline (Fig. 1A, B). It should be noted that ANO5 may play a role in Ca^2+^ homeostasis (8) and that this may contribute to the observed changes in the post-injury Ca^2+^ transients in ANO5-KO myofibers.Second, we employed a common damage assay based on the cell-impermeant dye FM1-43 (33, 39–41, 45, 52). FM1-43 is a water-soluble stearyl dye that is not fluorescent in aqueous solution, but fluoresces intensely when inserted into lipid-rich membranes. When the sarcolemma is damaged, FM1-43 enters the cell, labels internal membranes, and produces a bright fluorescent spot around the site of injury. If the membrane is quickly repaired, the quantity of dye entering the fiber is attenuated compared to fibers where repair is defective. In our system, FM1-43 fluorescence increases after injury about 3-times more rapidly in ANO5-KO muscle fibers than in WT. Fluorescence intensity doubled at t = 12 s and t = 36 s post-injury in ANO5-KO and WT fibers, respectively (Fig. 1C, D), similar to what has been previously reported under different experimental conditions (36). Comparison of the mean FM1-43 fluorescence revealed that dye accumulation was significantly greater in ANO5-KO fibers as soon as 8 s post-injury (Fig. 1D). Together, the data in Fig. 1 strongly support the idea that ANO5-deficient myofibers are defective in plasma membrane repair.

**Figure 1.**
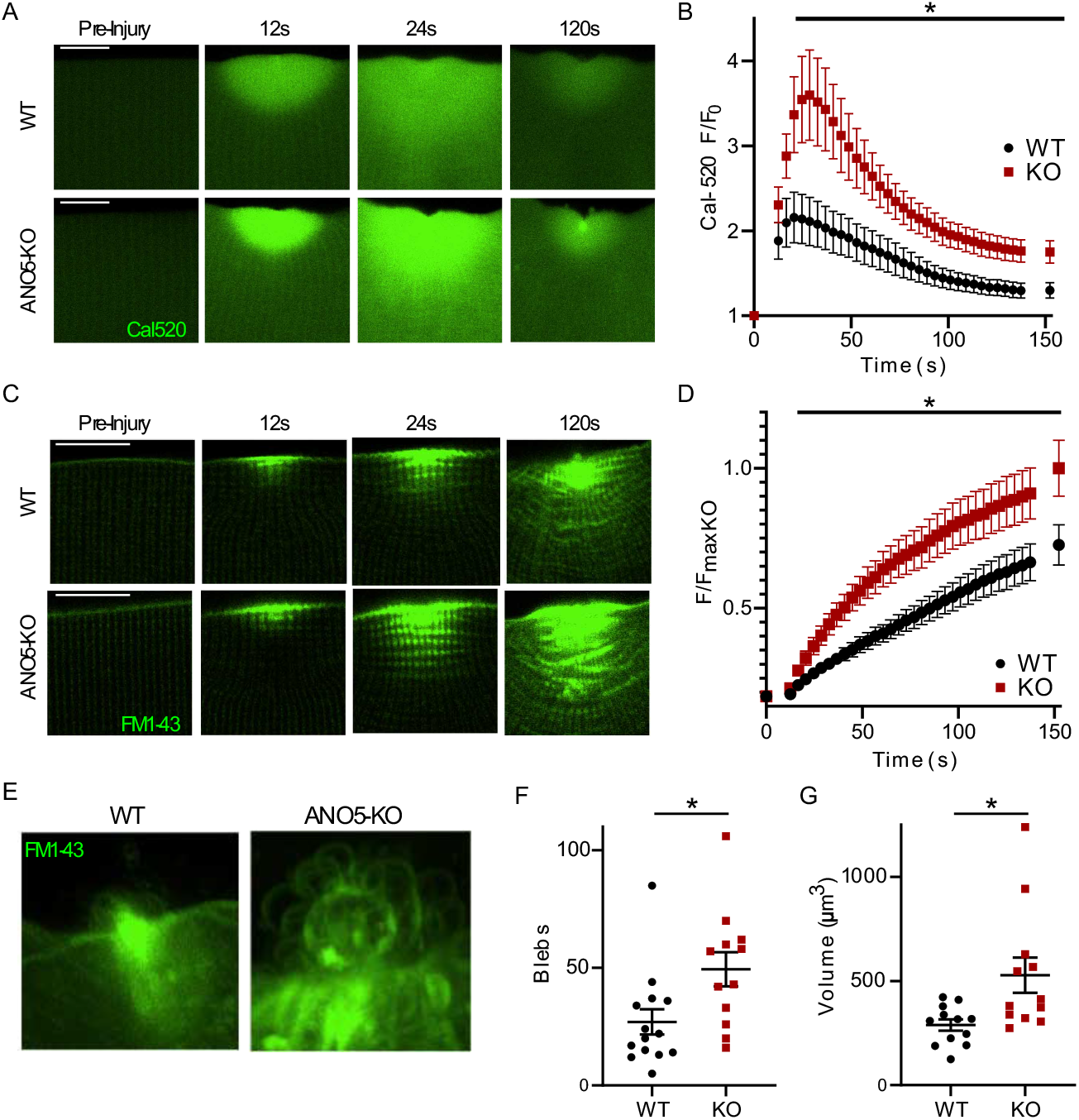
Plasma membrane repair is defective in ANO5-KO mouse muscle fibers. (A) Representative images of Ca^2+^ influx detected with the Ca^2+^-sensitive dye Cal-520 following laser-induced membrane damage in WT or ANO5-KO mice. Scale bar = 10 μm. (B) Time course of Cal-520 fluorescence intensity following membrane injury, normalized to initial fluorescence within the muscle fiber. Ca^2+^ is significantly higher in ANO5-KO fibers 20-150 s post-injury. ANO5-KO n = 7 fibers, WT n = 10 fibers. (C) Representative images of FM1-43 infiltration following membrane damage in WT or ANO5-KO mice. Scale bar = 10 μm. (D) FM1-43 intensity reported as fractional fluorescence F divided by average maximal fluorescence recorded in ANO5-KO myofibers (F_maxKO_). ANO5-KO n = 12, WT n = 15 fibers. FM1-43 dye accumulated significantly faster in ANO5-KO fibers beginning at 8 s post-injury and continued during 150 s of imaging. (E) Maximum intensity projections of FM1-43 dye generated from z-stacks of WT or ANO5-KO muscle fibers acquired approximately 10 min post-injury. Images with large differences in the x-y plane were chosen because information in the z-plane is not well rendered by maximum intensity projections. (F) Quantification of blebs (observed with FM1-43 dye) in the membrane-rich repair patch that forms following injury. ANO5-KO patches displayed significantly more membrane blebs than those from WT fibers. ANO5-KO n = 12 fibers, WT n = 14 fibers. (G) Quantification of patch volumes obtained by integrating patch areas from individual slices of z-stacks acquired approximately 10 min post-injury. Blebbing of the patch in ANO5-KO fibers contributed to a significant increase in overall patch volume. ANO5-KO n = 12 fibers, WT n = 12 fibers.

### The structure of the membrane repair patch is abnormal in the absence of ANO5

Sarcolemmal injuries in the presence of FM1-43 also revealed the formation of a large “cap” of fluorescence at the site of injury. This cap, which likely reflects processes invovled in the repair of the laser-induced lesion, was relatively small (compact) in WT fibers, but was large and heavily vesiculated in ANO5-KO fibers. While membrane blebs were occassionally observed in repairing WT myofibers as well, these blebs were seen to be released as extracellular vesicles (EVs) (Supplemental Movie 1), consistent with previous reports (53). The volume of FM1-43-positive membrane in the patch was significantly elevated in ANO5-KO fibers, which is plausibly explained by the retention of blebs/vesicles (Fig. 1E-G, Supplemental Movie 2). This difference in the structure of the repair cap and the difference in EV shedding is also revealed by the structure of the repair cap visualized by annexin accumulation as described in a subsequent section (see Fig. 4 and Supplemental Movies 4, 6, 7).

### ANO5 traffics to the site of injury

Given that muscle fibers lacking ANO5 are repair deficient, we sought to evaluate whether ANO5 localizes to the site of injury following plasma membrane damage. We electroporated plasmids encoding fluorescently-tagged ANO5 into muscle fibers of WT mice and performed laser injury. We observed fast accumulation of ANO5 at the plasma membrane adjacent to the site of damage (Fig. 2A, C), a region previously termed the wound “shoulder” (40). This behavior was recapitulated in ANO5-KO fibers, indicating that ANO5-KO myofibers retain the machinery necessary for ANO5 trafficking (Fig. 2A, C). The time course of accumulation of ANO5 was well fit by a single exponential with similar τ for WT and knockout fibers (WT τ = 19.9 s, ANO5 knockout τ = 17.5 s). This trafficking was not the result of local, injury-induced muscle contraction as sarcolemma-localized ANO1 did not accumulate to a comparable extent or with similar kinetics to ANO5 (Fig. S1). Since plasma membrane repair is Ca^2+^-dependent and because ANO5 is Ca^2+^-activated, we sought to characterize the kinetics of ANO5 trafficking to the wound shoulder with respect to the injury-induced Ca^2+^ transient. To that end, we co-electroporated ANO5-Neon with the genetically encoded Ca^2+^ sensor R-GECO1.2 (K_d_ = 1.2 μM) (54). As seen with Cal-520 (Fig. 1A), intracellular Ca^2+^ increased in a semicircular area immediately after injury (Supplemental Movie 3). ANO5 accumulation at the wound shoulder paralleled the time course of the Ca^2+^ transient. As the Ca^2+^ levels returned to baseline, ANO5 dispersed from the site of injury.

**Figure 2.**
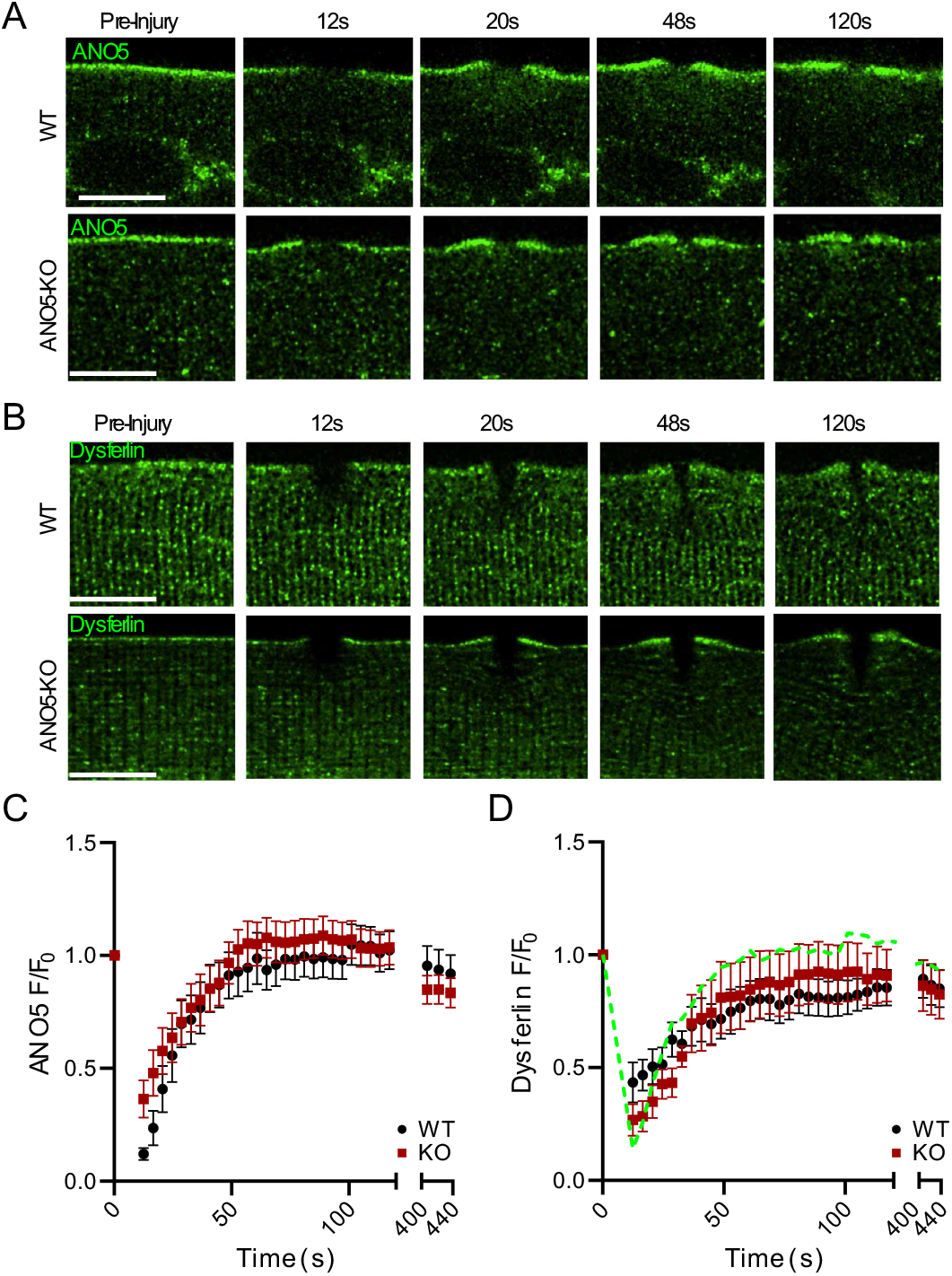
ANO5 accumulates at the plasma membrane in response to wounding. (A) Deconvolved images of ANO5 translocation to wound-adjacent plasma membrane in WT or ANO5-KO fibers at several time points after injury. Scale bar = 10 μm. (B) Deconvolved images showing dysferlin accumulation at wound-adjacent plasma membrane in WT or ANO5-KO fibers after injury, mirroring what is observed with fluorescently tagged ANO5. Scale bar = 10 μm. (C) Quantification of ANO5 fluorescence at the plasma membrane following injury in ANO5-KO (red squares) or WT (black circles) myofibers, normalized to initial plasma membrane fluorescence. ANO5-KO n = 16 fibers, WT n = 15 fibers. (D) Quantification of dysferlin fluorescence in ANO5-KO (red squares) or WT (black circles) myofibers following injury. The time course of hANO5 accumulation in WT fibers is shown as a green dotted line for reference. ANO5-KO n = 11 fibers, WT n = 9 fibers.

Overall, the timing and localization of ANO5 trafficking after wounding resemble that reported for dysferlin, a protein with well-established connections to plasma membrane repair (33, 38, 40, 43, 55). Dysferlin and ANO5 have been linked previously because of similarity in clinical manifestations of LGMDR2 and LGMDR12, but a relationship between the two in the repair pathway is not established. To test whether dysferlin trafficking is altered upon loss of ANO5, we examined the response of dysferlin-EGFP to laser damage in WT or ANO5-KO myofibers. The kinetics of dysferlin accumulation were nearly identical in WT (τ = 29.0) and ANO5-KO fibers (τ = 25.8) (Fig. 2B, D), though slower and less robust than ANO5. We conclude that the repair defect in ANO5-deficient muscle is independent of dysferlin.

### Annexins require ANO5 for normal trafficking

Annexins are phospholipid binding proteins with established roles in plasma membrane repair in various cell types (53, 56–59). Specifically in muscle, ANXA1, ANXA2, ANXA5, ANXA6 (mouse (38–41, 44)) and ANXA11 (zebrafish (42)) have been implicated in sarcolemmal resealing after injury. Annexins form a punctate “cap” that first appears within seconds of wounding, but continues to grow on a timescale of minutes (40, 42). Since our results (e.g. Fig. 1) indicate that resealing is delayed in ANO5-KO muscle fibers, we wanted to determine whether this could be attributed to abnormal development of the annexin-dependent repair cap. We expressed fluorescently tagged ANXA1, ANXA2, ANXA5 and ANXA6 (individually or together) in FDB fibers of WT or ANO5-KO mice and followed cap formation after injury (Fig. S2). All annexins tested trafficked to the cap with biphasic kinetics (Fig. 3, Table S2). To quantify the kinetics, F/F_0_ vs. time was fit to the sum of two exponentials (R^2^ > 0.995 for every curve). In WT, ANXA5 (τ_fast_ = 12.8, τ_slow_ = 142) and ANXA2 (τ_fast_ = 24, τ_slow_ = 118) accumulated most rapidly. By comparison, of the magnitude of ANXA6 accumulation was much larger, but proceeded over a longer period (τ_fast_ = 40.4, τ_slow_ = 830). ANXA2 and ANXA6 were affected dramatically by loss of ANO5, but in opposite directions. The amplitude of the slow component of ANXA2 accumulation was increased > 5-fold, corresponding to significantly greater overall amounts of ANXA2 in the repair cap at later timepoints (Fig. 3B, D, S2). In WT fibers ANXA2 was seen to be shed as extracellular vesicles from the cap in a manner like we described above for FM1-43 experiments (Supplemental Movie 4). Thus, ANXA2 may reach a steady state level during normal repair through opposing actions of trafficking and shedding, a balance dependent on ANO5. In contrast, the amplitude of both fast and slow components of ANXA6 were decreased ~2-fold by ANO5-KO. We conclude that loss of proper annexin trafficking is a key mechanism underlying deficient repair in ANO5-KO myofibers.

**Figure 3.**
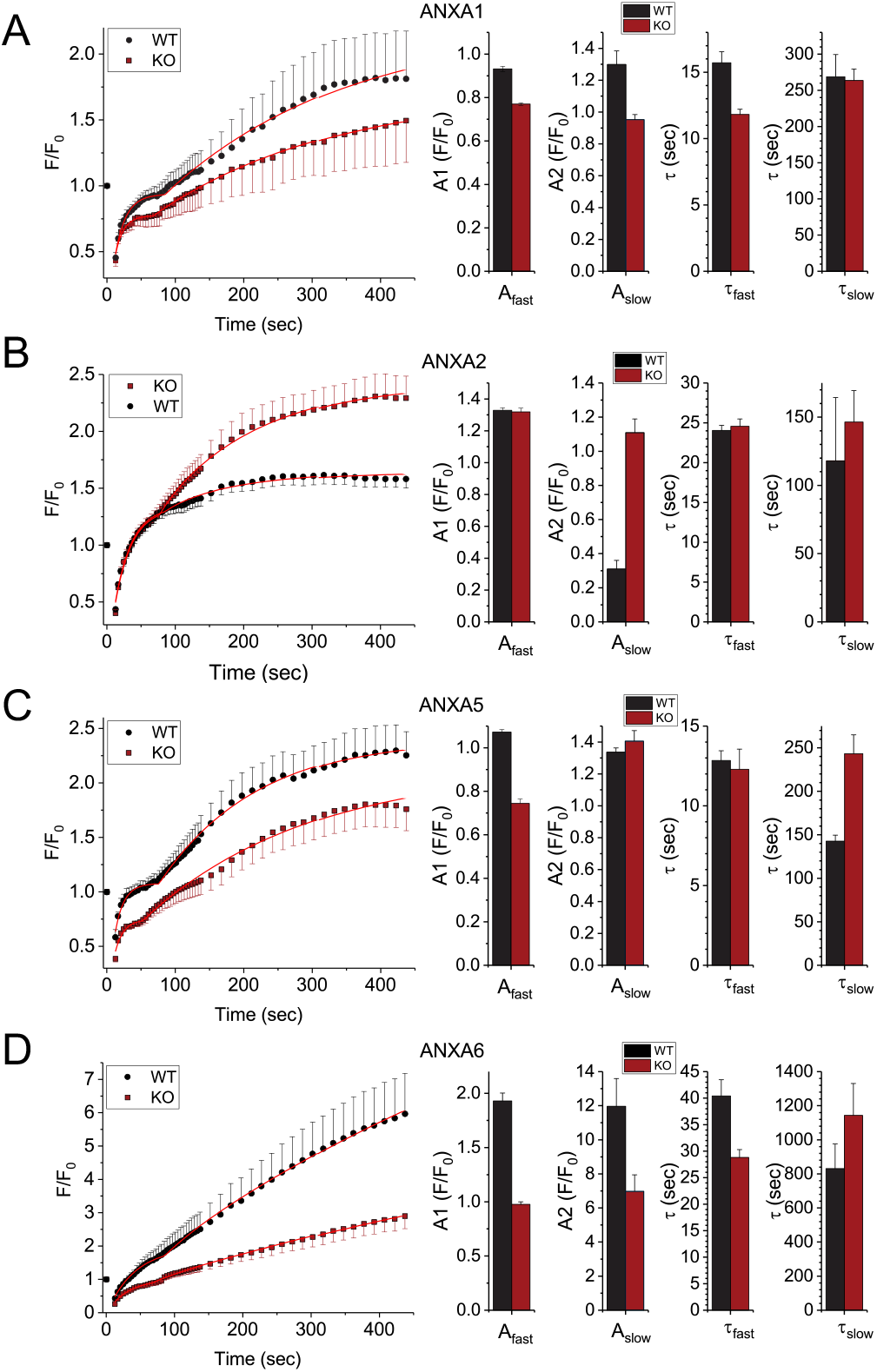
Kinetics of annexin trafficking are quantitatively altered with loss of ANO5. Time courses of annexin fluorescence were plotted as a function of time and fitted to the sum of two exponentials. Amplitudes, labeled A, and time constants, labeled τ, are shown for the fast (early) and slow (late) components of the curves. Raw and fitted data were compared for injured WT or ANO5-KO fibers expressing (A) ANXA1, (B) ANXA2, (C) ANXA5, and (D) ANXA6. ANXA1: ANO5-KO n = 13 fibers, WT n = 14 fibers; ANXA2: ANO5-KO n = 42 fibers, WT n = 37 fibers; ANXA5: ANO5-KO n = 36 fibers, WT n = 29 fibers; ANXA6: ANO5-KO n = 15 fibers, WT n = 18 fibers.

### Loss of ANO5 disrupts normal ANX repair cap architecture

When we co-expressed annexins in WT myofibers, we observed that different annexins occupied distinct spatial regions within the cap. To quantify this organization, ANXA1, ANXA5, and ANXA6 were each co-electroporated into fibers with ANXA2. After injury the intensity of each anenxin was measured relative to ANXA2 as described in Methods. These intensity values were normalized to maximal ANXA2 intensity and plotted vs distance from this maximal ANXA2 intensity. In WT fibers, ANXA5 was consistently concentrated towards the extracellular space relative to ANXA2, while ANXA6 and ANXA2 localizations largely overlapped. The opposite was observed in ANO5-KO fibers, where ANXA5 and ANXA2 appeared to colocalize, while ANXA6 was shifted toward the intracellular space relative to ANXA2 (Fig. 4A-D). For ANXA1, no differences were observed between WT and ANO5-KO fibers (not shown). Changes in the two-dimensional organization of the cap were accompanied by alterations in the volumes of the annexins within the cap. Orthogonal views, generated from z-stack images taken ~10 minutes after injuring fibers, revealed that ANXA2 extended a greater distance in all 3 spatial planes in ANO5-KO fibers than in WT fibers, and calculated ANXA2 repair cap volumes were significantly larger in ANO5-KO myofibers (Fig. 4E, F). In contrast, ANXA5 and ANXA6 caps were significantly smaller in ANO5-KO muscle (Fig. 4E, F). These results are in agreement with our findings that in ANO5-KO fibers, ANXA2 accumulates beyond normal physiological levels and ANXA5 and ANXA6 show depressed trafficking.

**Figure 4.**
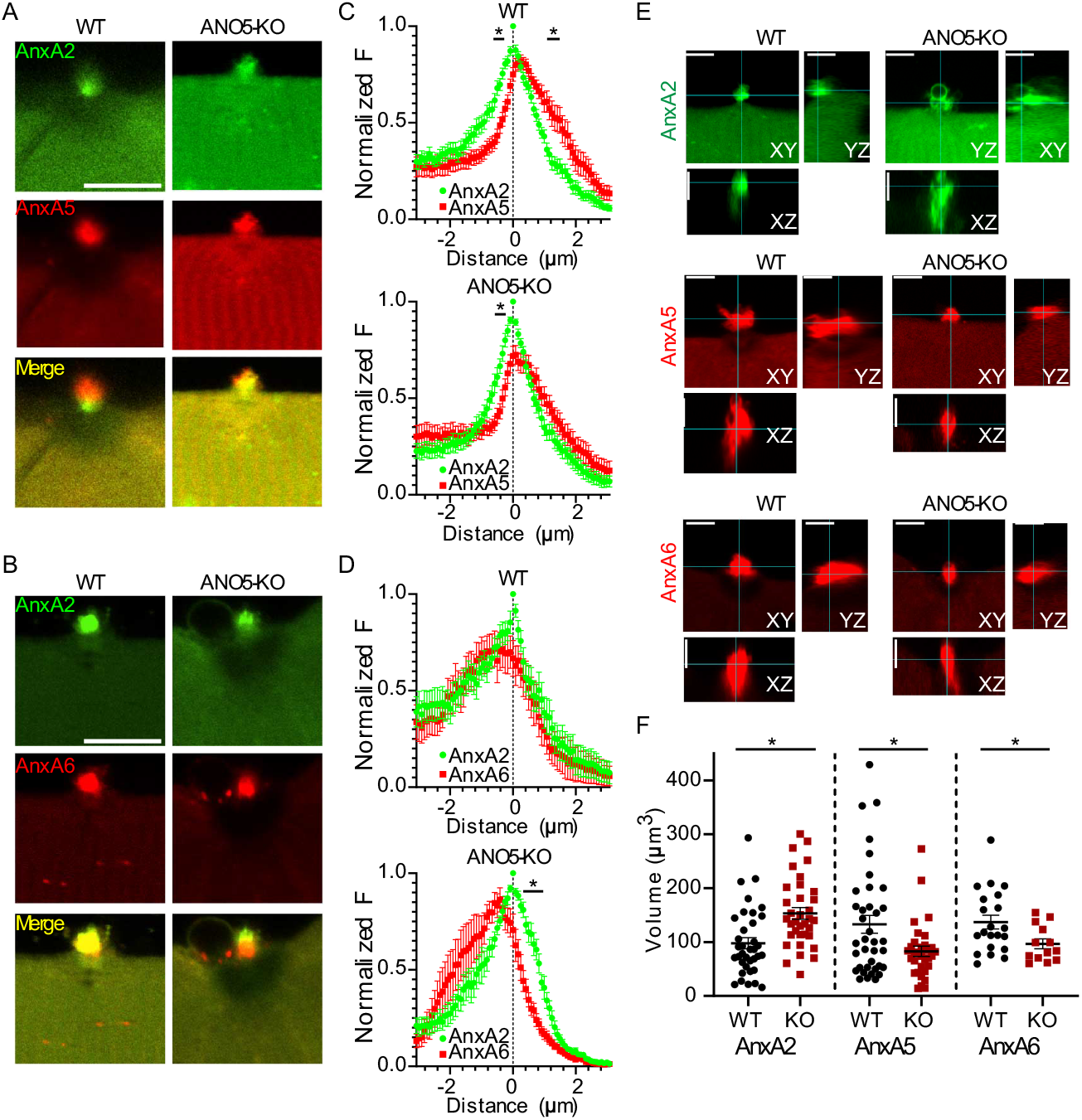
Architecture of the Annexin repair cap is altered upon loss of ANO5. Representative images showing localization of co-electroporated (A) ANXA2 and ANXA5 or (B) ANXA2 and ANXA6 in injured muscle fibers. ANXA2 and ANXA5 occupy overlapping but distinct spatial regions in WT fibers but appear in nearly identical regions of the repair cap in ANO5-KO fibers. In contrast, ANXA2 and ANXA6 show near complete overlap in WT fibers which is lost in the absence of ANO5. Scale bar = 10 μm. (C, D) Quantification of the spatial organization of Annexin proteins in the repair cap derived from line profiles drawn through the cap images taken approximately 7 min after injury. Maximum fluorescence values for ANXA2 are normalized to 1 and are defined as “distance” 0. Intensity was measured for co-electroporated annexins along the same line and compared to ANXA2. ANXA2 and ANXA5: ANO5-KO n = 23 fibers, WT n = 26 fibers; ANXA2 and ANXA6: ANO5-KO n = 9, WT n = 9 fibers. (E) Orthogonal views of the annexin repair cap generated from z-stacks acquired approximately 10 min post-injury. Scale bar = 5 μm. (F) Volume approximations made for individual Annexin proteins in repair caps following injury. Z-plane images were taken every 0.5 μm, and volumes were estimated by summing products of the area of the cap in a given z-image by the depth of the z-slice. ANXA2 caps were significantly larger in injured ANO5-KO fibers, while ANXA5 and ANXA6 caps were significantly smaller. ANXA2: ANO5-KO n = 35 fibers, WT n = 36 fibers. ANXA5: ANO5-KO n = 32 fibers, WT n = 37 fibers. ANXA6: ANO5-KO n = 13 fibers, WT n = 21 fibers. In all panels: *, p < 0.05.

### PtdSer and Phosphatidylethanolamine (PtdEtn) are exposed after injury

The inner leaflet of the plasma membrane is normally enriched in PtdSer and PtdEtn. Phospholipid scrambling results in the exposure of these phospholipids to the extracellular face of the membrane. Accumulation of PtdSer at the site of plasma membrane injury has been documented, but the mechanism by which PtdSer comes to be exposed extracelluarly during repair is unresolved (40, 43). We previously showed that ANO5 supports Ca^2+^-dependent PLS (21). Therefore, we hypothesized that PtdSer exposure during plasma membrane repair is ANO5-dependent. To test this, we laser-injured WT or ANO5-KO fibers in the presence of the PtdSer-binding C2 domain of Lactadherin fused to Clover or mCherry fluorescent proteins (LactC2-FP). Contrary to our expectations, LactC2-FP accumulated at the site of injury rapidly and robustly in both WT and ANO5-KO fibers, indicating that ANO5 is not required for accumulation of PtdSer in membrane patches (Fig 5A, B). Although ANO scramblases are generally considered non-specific with regard to their phospholipid substrates (60, 61), we wanted to examine whether loss of ANO5 reduces exposure of lipids besides PtdSer. We conducted injury experiments in the presence of Cy3-conjugated duramycin, which binds specifically to PtdEtn. Duramycin-Cy3 accumulated rapidly at the patch site, with localization comparable to that observed with LactC2 (Fig 5C, D). Surprisingly, although PtdEtn accumulation was similar at the end of the 440 s recording period in WT and ANO5-KO fibers, initial PtdEtn accumulation was significantly less in ANO5-KO fibers. At 100 s, PtdEtn accumulation was ~44% less in ANO5-KO compared to WT. Although the results of Fig. 5D could be interpreted as supporting the idea that ANO5 selectively scrambles PtdEtn, additional experiments in the next section show that ANO5-dependent scrambling is not required for normal membrane resealing. Furthermore, it is unlikely that lipid exposure in ANO5-knockout fibers can be explained by compensatory ANO6 expression (and consequent upregulation of ANO6-PLS), because ANO6 levels are unchanged in ANO5-KO myocytes and myotubes relative to WT (Fig. S3, (21)). We also observed PtdSer and PtdEtn exposure even in fibers injured in Ca^2+^-free solution, suggesting that PtdSer and PtdEtn exposure is not dependent on Ca^2+^ influx (Supplemental Movie 5). Together, these findings seeem to rule out ANO5 and ANO6 as primary mediators of PtdSer exposure during membrane repair and suggests that lipid probe accumulation in our system reflects membrane trafficking at the damage site. If that is true, reduced PtdEtn exposure in the initial phase of repair is consistent with a decline in accumulation of membranes/intracellular vesicles utilized for wound sealing.

**Figure 5.**
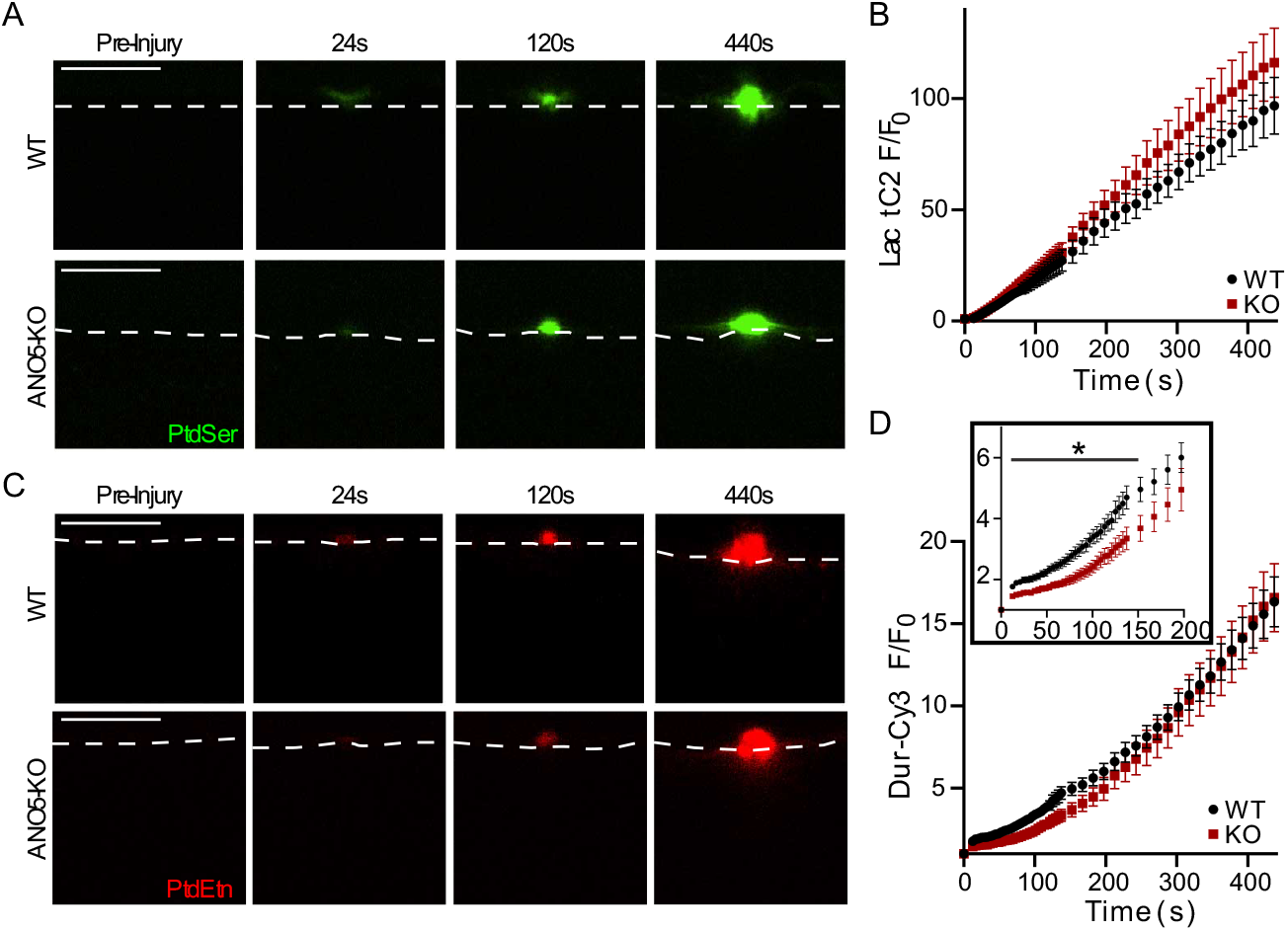
ANO5 is not required for exposure of intracellular lipids following plasma membrane injury. (A) Representative images showing accumulation of the fluorescent PtdSer-sensor LactC2-Clover after laser-mediated plasma membrane injury. The cell boundary is indicated by a dotted white line. Scale bar = 10 μm. (B) Time course of LactC2-FP (indicating PtdSer) accumulation after damage, normalized to initial fluorescence at the injury site. PtdSer accumulation at the wound is not impaired in the absence of ANO5. ANO5-KO n = 23 fibers, WT n = 14 fibers. (C) PtdEtn, detected by Cy3-conjugated duramycin, appears rapidly at the repair patch following wounding. The cell boundary is marked by a dotted white line. Scale bar = 10 μm. (D) Quantification of PtnEtn kinetics, indicating rapid, ANO5-independent recruitment of PtdEtn to the extracellular surface of muscle fibers after damage. PtdEtn accumulation was greater in WT up to 152s, but then equalized between groups. Inset in top left of plot highlights the difference in initial PtdEtn accumulation. ANO5-KO n = 9 fibers, WT n = 21 fibers.

### ANO5 “scrambling domain” is not required for repair

We and others have identified a conserved 34 amino acid region of ANO5 necessary for ANO5-mediated PLS and ionic currents (17, 21). Replacement of this scrambling domain with the corresponding sequence from the chloride channel ANO1 to form chimeric “ANO5-1-5”, destroys ANO5-dependent phospholipid scrambling (21). Given that PtdSer and PtdEtn are exposed during membrane repair even in the absence of ANO5, we hypothesized that muscle membrane repair proceeds independently of ANO5-dependent PLS. To test this, we electroporated fluorescently-tagged ANO5 or ANO5-1-5 into ANO5-KO myofibers and performed laser damage in the presence of FM1-43. Generally, the expression of ANO5-1-5 was low, and tag-associated fluorescence was weak and diffuse. Nevertheless, both ANO5 and ANO5-1-5 significantly reduced FM1-43 infiltration post-injury (> 2-fold increase in the time of FM1-43 fluorescence doubling over F_0_ relative to ANO5-KO alone) (Fig. 6A, C). Consistent with a “rescue” effect of ANO5 and ANO5-1-5, the FM1-43-positive cap in ANO5-KO fibers expressing either construct was significantly reduced compared to ANO5-KO alone (Fig. 6D). To test whether these ANO5 constructs promote repair through restoration of normal annexin trafficking, we co-expressed ANO5 or ANO5-1-5 with ANXA2 or ANXA6. ANO5 overexpression reduced ANXA2 accumulation at all timepoints, yielding an F/F_0_ value 106% of WT (68% of ANO5-KO) at the conclusion of the experiment. Interestingly, ANO5-1-5 did not reduce ANXA2 accumulation in ANO5-KO fibers (final F/F_0_ was 159% of WT (102% KO)) (Fig. 6E and Supplemental Movie 6). In contrast, both ANO5 and ANO5-1-5 markedly improved ANXA6 trafficking to the lesion in ANO5-KO fibers. ANO5 restored final ANXA6 F/F_0_ to 72% of WT (149% of KO), while ANO5-1-5 improved ANXA6 F/F_0_ to 88% of WT (182% KO) (Fig. 6F and Supplemental Movie 7). Because ANO5-1-5 expression reverses ANXA6 but not ANXA2 phenotypes while also substantially reducing FM1-43 accumulation in ANO5-KO myofibers, it appears likely that ANXA6 translocation to the repair cap is crucial for efficient wound resealing. However, our results suggest that the ANO5 scrambling domain enables release of ANXA2-positive EVs, which may have a signaling role during repair *in vivo* (62).

**Figure 6.**
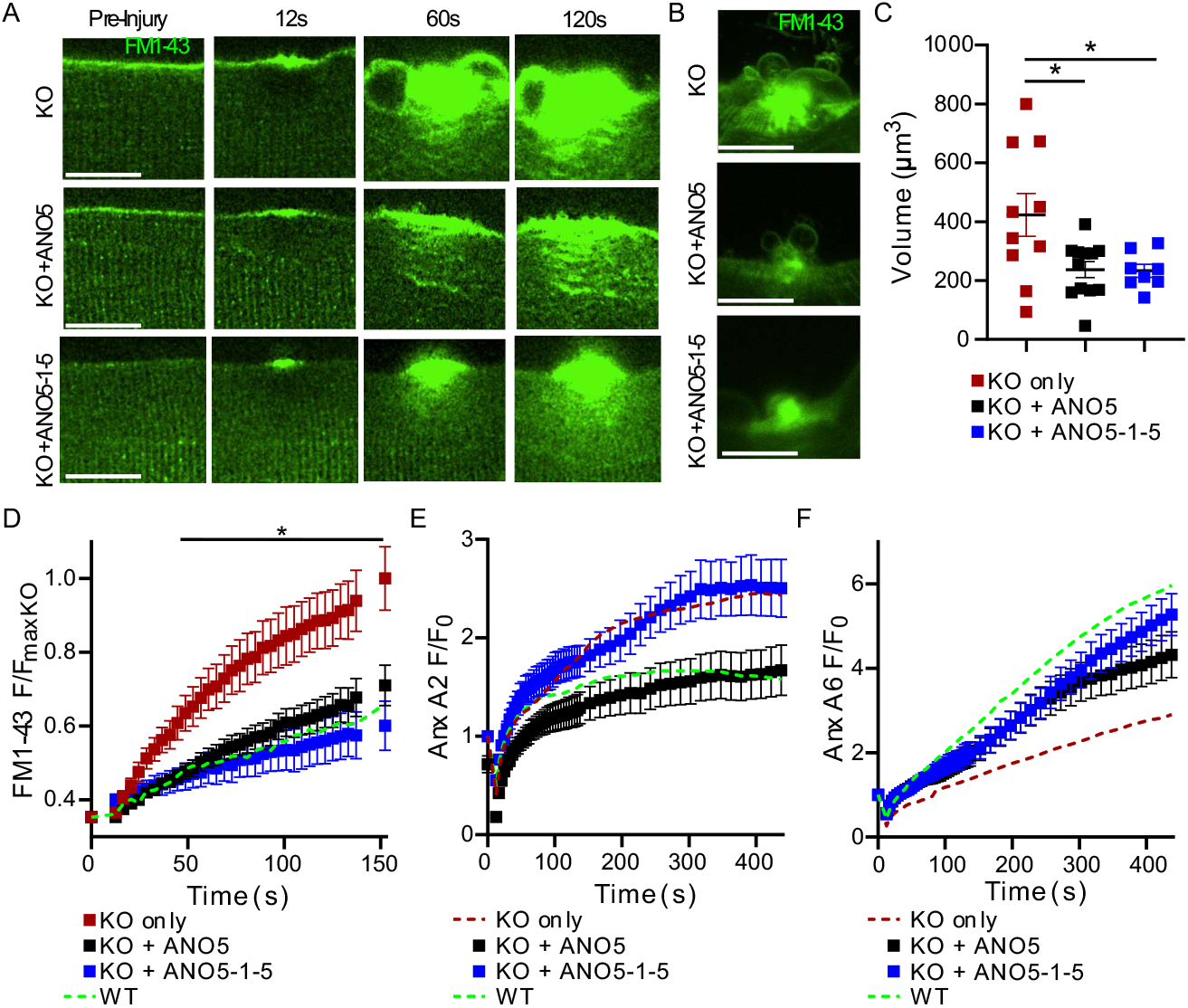
ANO5 does not require “scrambling domain” to rescue defective repair. (A) Deconvolved images of FM1-43 dye infiltration in ANO5-KO fibers alone (top), ANO5-KO fibers expressing WT hANO5 (middle), or ANO5-KO fibers expressing mutant ANO5-1-5 lacking PLS function, prior to and following laser-induced injury. Scale bar = 10 μm. (B) Maximum intensity projections of z-plane image stacks from ANO5-KO fibers, ANO5-KO fibers expressing WT hANO5, and ANO5-KO fibers expressing mutant ANO5-1-5 (top, middle, bottom, respectively) acquired 10 min post injury. Scale bar = 10 μm. Images with large differences in the x-y plane were chosen because information in the z-plane is not well rendered by maximum intensity projections. (C) Quantification of FM1-43-demarcated patch volumes obtained by integrating patch areas from individual slices (separated by 0.5 μm) of z-stacks acquired ~10 min post-injury. ANO5-KO n = 10 fibers, ANO5-KO + ANO5 n = 12 fibers, ANO5-KO +ANO5-1-5 n = 8 fibers. (D) Quantification of FM1-43 accumulation in ANO5-KO fibers and ANO5-KO fibers that have been “rescued” with WT hANO5 or chimeric ANO5-1-5. Data are reported as fractional fluorescence F divided by average maximal fluorescence recorded in ANO5-KO myofibers (F_maxKO_). FM1-43 accumulation was significantly reduced in fibers expressing WT hANO5 (open squares, starting at 20s) and ANO5-1-5 (black filled squares, starting at 32s). ANO5-KO n = 11 fibers, ANO5-KO + hANO5 n = 16 fibers, ANO5-KO + ANO5-1-5 n = 10 fibers. A green dotted line represents the average FM1-43 F/F_maxKO_ value from 11 injured WT fibers as a reference. (E) ANXA2 accumulation, normalized to initial fluorescence, in ANO5-KO fibers expressing WT hANO5 (black squares) or chimeric ANO5-1-5 (blue squares). Green and red dotted lines representing ANXA2 accumulation in WT and ANO5-KO fibers, respectively, are shown as a reference. KO +ANO5 n = 32 fibers, KO +ANO5-1-5 n = 16 fibers. (F) ANXA6 accumulation, normalized to initial fluorescence, in ANO5-KO fibers expressing WT hANO5 (black squares) or chimeric ANO5-1-5 (blue squares). Green and red dotted lines representing ANXA2 accumulation in WT and ANO5-KO fibers, respectively, are shown as a reference. KO + ANO5 n = 24 fibers, KO + ANO5-1-5 n = 18 fibers. Data are presented as mean ± S.E.M. *, p < 0.05.

### An R58W mutation found in human patients produces defective repair in human myotubes

To determine whether membrane repair is also defective in human patients with ANO5 mutations, we isolated MPCs from a human patient carrying a homozygous c.172C>T (p.R58W) point mutation. This variant is classified as likely to be pathogenic (https://www.ncbi.nlm.nih.gov/clinvar/) because it is consistent with an autosomal recessive inheritance pattern of LGMDR12 and is predicted to disrupt ANO5 secondary structure (63–67). Differentiated patient myotubes laser damaged in the presence of FM1-43 showed a markedly elevated uptake of FM1-43 (indicating a poor repair response) relative to healthy control myotubes. Furthermore, FM1-43 accumulation was accompanied by significant plasma membrane blebbing in patient cells (Fig. 7A, B). After 2 minutes, FM1-43 fluorescence was 2.5-times greater in the R58W cells than in control cells (Fig. 7B). We reasoned that if deficient repair in patient cells were caused by ANO5 loss of function associated with the R58W mutation, this variant ought to be incapable of rescuing repair in ANO5-KO mouse myofibers. We electroporated ANO5-R58W-mCherry into ANO5-KO fibers and found that, at least under condition of overexpression, it was well expressed (Fig. S4), in contrast to the previously reported R758C patient mutation, which leads to ANO5 degradation (8). Expression of ANO5-R58W in ANO5-KO fibers had no effect on FM1-43 accumulation or patch volumes (Fig. 7C-E). We therefore conclude that pathogenicity associated with ANO5-R58W is related to plasma membrane repair incompetence.

**Figure 7.**
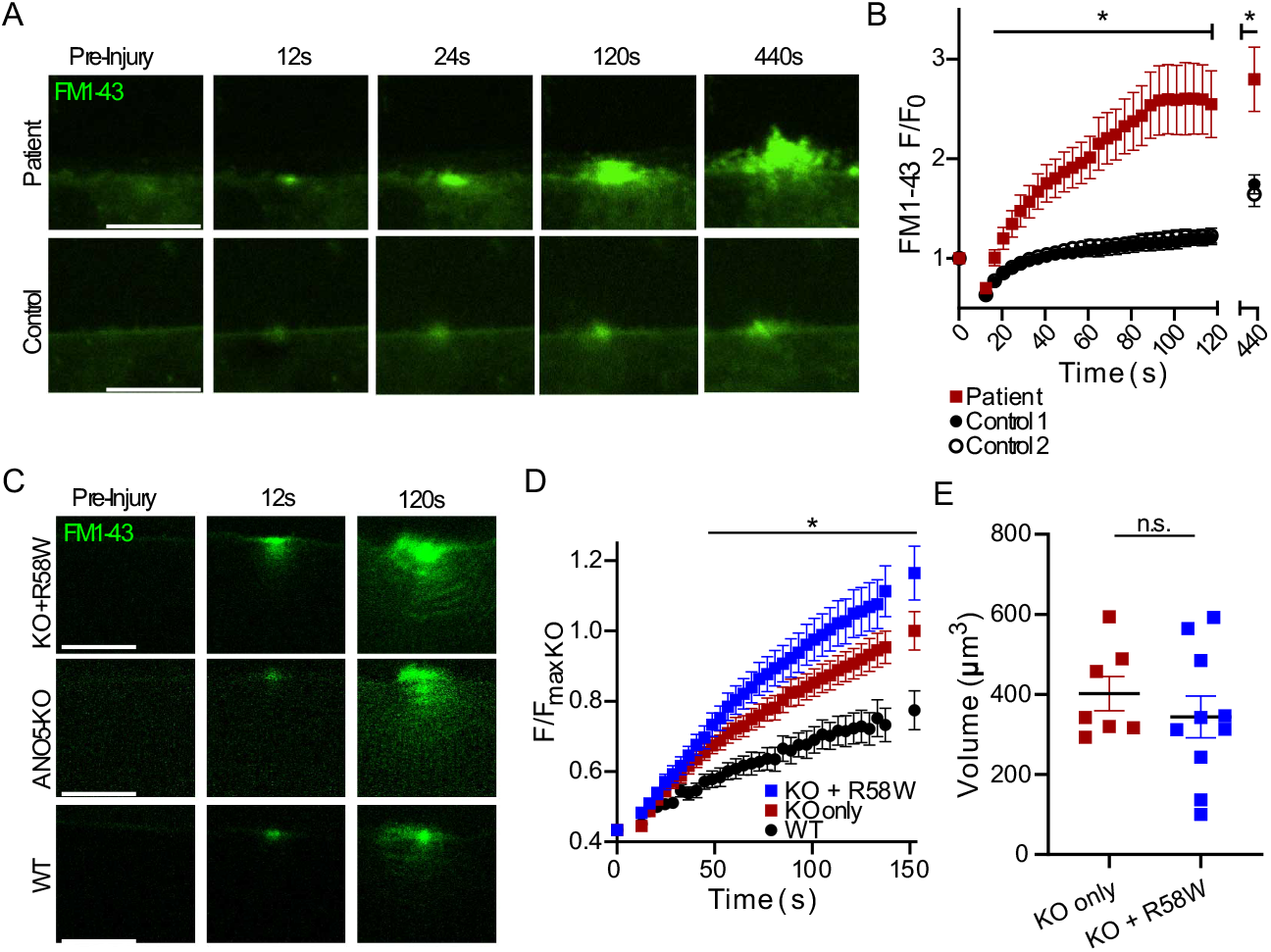
An R58W mutation found in human patients causes defective repair in cultured myotubes. (A) Representative images of FM1-43 infiltration in control or R58W patient human myocytes following laser-induced injury. (B) Quantification of FM1-43 accumulation following injury, which was significantly increased in R58W myocytes relative to controls. Cells derived from two healthy control individuals were used. Control 1 n = 9 cells, Control 2 n = 8 cells, Patient n = 11 cells. (C) Images showing FM1-43 uptake in injured mouse ANO5-KO, ANO5-KO expressing mutant R58W ANO5, and WT myofibers. (D) Kinetics of FM1-43 accumulation in mouse myofibers with or without expression of R58W ANO5. R58W ANO5 does not improve repair in ANO5-KO fibers, as dye accumulation in both ANO5-KO and ANO5-KO + R58W fibers was significantly greater than in WT. Data are reported as fractional fluorescence F divided by average maximal fluorescence recorded in ANO5-KO myofibers (F_maxKO_). ANO5-KO n = 17 fibers, ANO5-KO +R58W n = 25 fibers, WT n = 11 fibers. (E) Quantification of approximate FM1-43 demarcated patch volumes obtained from integrating patch areas from individual slices (separated by 0.5 μm) of z-stacks acquired approximately 10 min post-injury. ANO5-KO n = 7 fibers, ANO5-KO +R58W n = 10 fibers. All data are presented as mean ± S.E.M. *, p < 0.05.

## Discussion

### A role for ANO5 in membrane repair

Our results provide additional support for the growing literature implicating ANO5 in muscle membrane repair while providing novel mechanistic insights about this process. Consistent with previous results (8, 52), we show that ANO5-KO myofibers are more permeant to FM1-43 dye after injury. It is important to note that the FM1-43 assay, by itself, is incapable of resolving whether fibers are defective in repair or more susceptible to injury. Similarly, our finding that Ca^2+^ transients are altered in the absence of ANO5 cannot conclusively distinguish between defective repair and increased fragility. The matter is further complicated by the fact that, at least in myoblasts, it has been reported that ANO5 regulates Ca^2+^ homeostasis (8, 68). Chandra et al. demonstrated that injury-induced Ca^2+^ transients were prolonged in myoblasts from a LGMDR12 patient with the R758C mutation. They concluded that these prolonged transients were explained by defective uptake of Ca^2+^ into the ER because the slower Ca^2+^ transients correlated with ER fragmentation after injury (8). However, it is not known whether this feature is specific to the R758C mutation or whether ER fragmentation and slow Ca^2+^ transients are causally related. While chelation of Ca^2+^ was sufficient to reduce FM1-43 infiltration in immortalized LGMDR12 patient myoblasts, it does not rule out the possibility that a reduction in free intracellular Ca^2+^ simply reduces cellular injury. In ANO5 knockout muscle fibers, we observed an increase in the amplitude of the Ca^2+^ transient and a much more pronounced slowing of its return to baseline. Ca^2+^ indicator fluorescence represents the difference between Ca^2+^ influx, which is dependent both on the size of the wound and the rate at which it is repaired, and the removal of Ca^2+^ through uptake into cellular organelles or extrusion from the cell. Larger initial Ca^2+^ amplitude is consistent with increased damage and/or reduced repair, while slower recovery of intracellular Ca^2+^ is consistent with delayed repair and/or defective Ca^2+^ uptake or efflux.

We found that ANO5 rapidly condensed along the plasma membrane immediately next to laser-induced lesions (Fig. 2A, C). We suggest that this is an active process because previous studies using plasma membrane targeted fluorescent proteins in zebrafish muscle did not find evidence of non-specific enrichment of plasma membrane proteins at wounds (42, 43). Furthermore, we show that sarcolemma-localized ANO1 does not accumulate at wound sites to the same extent that ANO5 does (Fig. S1). Because accumulation of ANO5 at wounds was coincident with Ca^2+^ influx (Supplemental Movie 3) that caused visible local contraction, the possibility exists that localized contraction of cortical actin might be involved in drawing plasma membrane proteins towards the wound (69, 70). Interestingly, both dysferlin (69) and annexins (40) translocate to wounds actin-dependently. Regardless of its mechanism of trafficking, the appearance of ANO5 at wound shoulders likely places it at a favorable location to facilitate formation of the annexin cap to reseal damaged myofibers. Additional work is required to clarify whether ANO5 mediates annexin cap formation through direct interaction of annexins, or if it is required at another, upstream step in myofiber repair. However, the recent finding that the ANO5 paralog ANO6 is involved in nonconventional annexin secretion (71) favors such a model.

### ANO5-dependent lipid sorting is not required for muscle membrane repair

Phospholipids are emerging as central players in membrane repair processes. Previous studies have demonstrated rapid accumulation of PtdSer and phosphatidylinositol (4,5)-bisphosphate at or adjacent to damage sites in muscle. However, reports vary with respect to precisely which lipids are required to facilitate repair and where they are needed. For example, Demonbreun et al. (40) show that PtdSer accumulated at the shoulder of the wound in mouse myofibers. These studies employed genetically encoded, intracellularly expressed LactC2-GFP as a PtdSer probe. This approach has disadvantages because the probe is likely to bind to PtdSer prior to the injury and this could interfere with downstream events. Furthermore, it does not provide an unambiguous conclusion regarding whether phospholipid scrambling is involved because it cannot distinguish between PtdSer located in the inner and outer membrane leaflets. Middel et al. (43) attempted to mitigate this problem by expressing a secreted form of Annexin-V-YFP and found that PtdSer localized to the repair patch itself, rather than the wound shoulder. This result agrees with our finding using a different approach: prior to injury we bathed fibers in purified LactC2-FP protein (72). Because LactC2 protein is impermeant, it labels only PtdSer in the external leaflet of the membrane, although it is uncertain whether PtdSer exposure occurs as a result of phospholipid scrambling or some other mechanism, such as vesicular fusion.

We were surprised to find that LactC2-FP and duramycin-Cy3 accumulation in both WT and ANO5-KO fibers was robust after injury. Accumulation of these probes was restricted to the repair cap itself rather than the shoulder of the wound. Also, we observed that of the accumulation of these probes at damage sites occurred in the absence of extracellular Ca^2+^ (Supplemental Movie 5). This suggests the possibility that acidic phospholipid exposure may be mediated by MG53-mediated intracellular vesicle aggregation (49) and the Ca^2+^-independent interaction between the arginine-rich C-terminus of dysferlin and PtdSer (43). Our results exclude ANO5-PLS as a repair mechanism in isolated myofibers and further solidify the distinct roles of dysferlin and ANO5 in membrane repair. Physiologically, the exposure of PtdEtn may be important because different annexins exhibit differential binding to different phospholipid species. For example, only ANXA2 and ANXA6 bind to PtdEtn-containing vesicles (73) and the binding of ANXA5 to PtdSer vesicles is modulated by PtdEtn (74). It is paradoxical that PtdSer exposure does not occur in patch-clamped cultured myogenic precursor cells from ANO5-KO mice (21) while PtdSer exposure triggered by membrane damage is apparently intact in adult ANO5-KO muscle fibers. This may be explained by differences (i) between myogenic precursor cells and adult muscle fibers or (ii) the spatio-temporal features or magnitude of the Ca^2+^ signal. Regardless, PtdSer exposure in damaged aldult fibers appears to be ANO5-independent.

### ANO5 and dysferlin function separately

Several annexin proteins have been linked to myofiber repair, through their presence in a punctate cap that develops in response to laser wounding (39–41, 44, 45). ANXA1 and ANXA2 reportedly interact with dysferlin to mediate repair in cultured myoblasts/myotubes (38), while a trafficking-deficient ANXA6 mutant inhibits dysferlin accumulation in wounded myofibers (40). This would seem to place dysferlin parallel to - or downstream of - ANXA6 in the repair pathway. Yet, in ANO5-KO mouse myofibers, where ANXA6 trafficking is depressed (Fig. 3D, S2), dysferlin behavior is apparently unchanged (Fig. 2B, D). Similarly, in zebrafish lacking ANXA6, dysferlin was unaffected (42). Thus, dysferlin is probably not dependent *per se* on ANXA6 for its translocation, but the two proteins might share a common subcellular compartment at some step during repair so that certain ANXA6 mutants are capable of preventing progression of dysferlin to the lesion. Additionally, it likely indicates that repair in muscle may not be a single process so much as several parallel and overlapping processes working in concert to effectively reseal the sarcolemma. We must also note that we observed somewhat limited dysferlin trafficking in WT fibers, which seems to contrast with previous reports (33, 69). Since the dysferlin C-terminus (42, 43, 75), rather than full-length dysferlin, seems to accumulate during repair, one possibility is that only some dysferlin expressed in our system is accessible to proteolytic cleavage. Absent a definitive explanation, it is nevertheless likely that ANO5 exerts minimal influence on dysferlin during the repair; whether the converse is true is an interesting point for future study.

### A coordinated annexin response reseals damaged myofibers

Given that different annexins interact differently with membranes to induce curvature (76, 77), it is important to appreciate the fine structure of the annexin cap in order to clarify how annexins facilitate wound healing. We observed a number of deficits with respect to the trafficking and final localization of ANXA2, ANXA5 and ANXA6, which together reveal the complexity of the wound response in muscle fibers. First, we noted delayed ANXA5 acummulation in the absence of ANO5 (Fig. 3C, S2). ANXA5 is capable of homo-oligomerizing to form two-dimensional arrays, which are associated with membrane resealing (44, 45). We also found that the relative positioning of ANXA5 and ANXA2 were different in WT and ANO5-KO fibers. One possiblity is that ANXA5 forms an initial proteolipid “scab” away from the fiber while additional membrane/protein is trafficked to the wound. ANXA2 has been shown to foster membrane blebbing, presumably through binding to the outer surface of membrane lesions, while both ANXA2 and ANXA6 are capable of inducing “membrane folding,” which involves binding and pulling together membranes. ANXA6 in particular has been suggested as a key factor in collapsing membrane lesions because it has two annexin cores and can consequently bind two membranes (41, 76). We identified depressed trafficking of ANXA6 overall in ANO5-KO myofibers, which is consistent with delayed plasma membrane resealing. In addition, whereas ANXA2 translocation to the repair cap plateaued within a few minutes after wounding in WT fibers, it continued to accumulate at the lesion in ANO5-KO fibers. ANXA2 is a driver of membrane bleb formation and marks blebs in the repair cap of ANXA6-deficient muscle (42). Thus, one intriguing possibility is that prolonged accumulation of ANXA2 in ANO5-KO fibers may be a factor in increased blebbing of the repair patch during FM1-43 experiments (Fig. 8). While altered annexin trafficking might be explained as an effect, rather than a cause, of defective repair in ANO5-KO fibers, our finding that ANO5-1-5 was capable of rescuing repair (according to FM1-43 infiltration) without restoring normal ANXA2 trafficking stands as a counterpoint to this idea.

**Figure 8.**
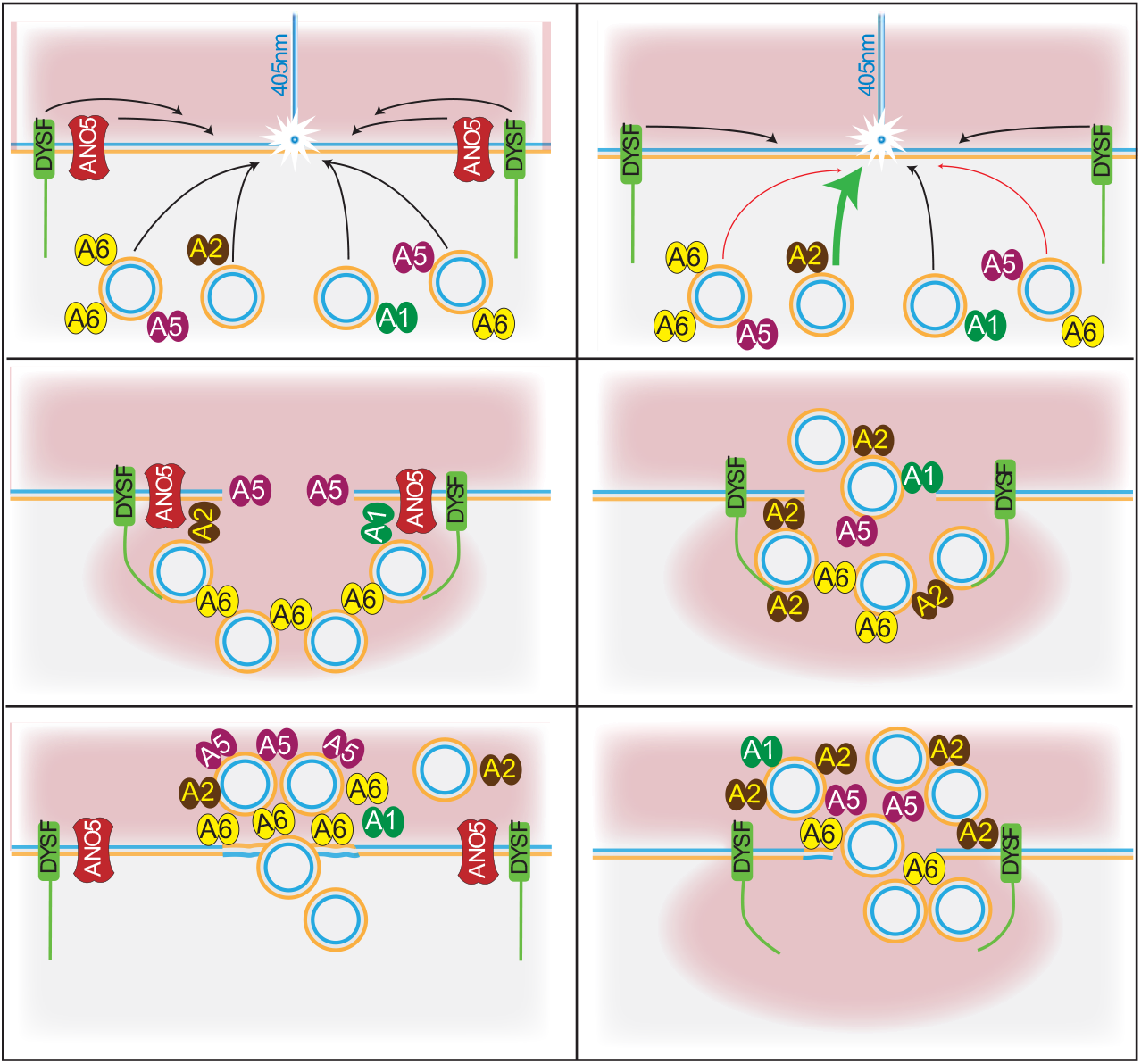
ANO5 coordinates annexin-mediated plasma membrane resealing. *Left.* Repair response in WT myofibers. Top panel: Dysferlin and ANO5 move laterally along the plasma membrane to facilitate proper organization of the annexin repair cap. Middle panel: Ca^2+^ (red) enters the fiber, initiating the repair cascade. ANXA5 is rapidly moved outside fiber, where it may form an initial proteolipid barrier as additional protein/membrane is trafficked for wound closure (middle panel). Bottom panel: ANXA6 binds phospholipids on multiple membrane surfaces to enable resealing. ANXA2 accumulates at the wound but is also shed in vesicles. *Right.* Repair is altered in ANO5-KO fibers. Top panel: Dysferlin translocates to the wound shoulder after damage, but ANX responses are abnormal in the absence of ANO5. Middle panel: ANXA5 traffics slowly and does not form a barrier. ANXA6 repair response is muted, while ANXA2 becomes over-represented in the repair cap. Bottom panel: Overabundance of ANXA2 leads to blebbing of the repair cap, but vesicles are not released. The repair cap is disorganized and resealing is delayed, leading to prolonged elevation of intracellular Ca^2+^.

The ANO5 paralog ANO6 has recently been proposed to play a role in repair of membranes damaged by bacterial toxins (78). Furthermore, ANO6-dependent scrambling of membrane lipids has been reported to be involved in translocation of ANXA2 and ANXA5 from the cytofacial surface of the plasma membrane to the exofacial surface in HeLa cells (71). This strengthens the link between ANOs, annexins, and membrane repair, but more importantly it raises the possibility that ANO5 and ANO6 could perform similar (though separate) functions in muscle. For example, ANO5 and ANO6 may have distinct annexin specificities and coordinate the annexin response during repair cooperatively by faciliating transport of particular annexin proteins to the wound. Loss of either ANO could result in a misbalance of annexin species within the repair cap, thus leading to a reduction in sarcolemmal resealing similar to what we have described here.

### What is the cell biology of ANO5?

ANO5 is most closely-related to ANO6, which is a well-characterized PLSase (19, 79, 80). While ANO6-mediated scrambling has an important role in blood clotting (e.g. (19)), and the scramblase domain of ANO6 is highly conserved in ANO5 (21, 80), a physiological role for ANO5-mediated scrambling has not been unambigiously demonstrated. We previously showed that ANO5-KO MPCs are PLS-deficient, and while this can be plausibly linked to a deficit in cell-cell fusion, it has not been conclusively connected (21, 36). Inhibition of ANO-mediated PtdSer exposure with the inhibitor CaCCinh-A01 reduced fusion of osteoclast precursors (84), providing more direct evidence for ANO scrambling in cell-cell fusion. Here, we provide evidence that ANO5 PLS is not required for efficient myofiber patch repair. We also show that LGMDR12 human patient myotubes carrying a homozygous R58W mutation are repair-deficient, but it is still unknown whether defective membrane repair is a common feature of all ANO5 mutations. The only other ANO5 patient mutation tested for defective membrane repair is an R758C missense mutation, which leads to ANO5 protein degradation (8). Furthermore, this defect was investigated only in myoblasts, rather than differentiated muscle cells, where ANO5 levels are highest (Fig. S3) and there are likely to be important differences in the repair pathway (42, 45). Overall, 72 ANO5 mutations in ClinVar have been described by at least one source as likely pathogenic or pathogenic, yet only 4 of these reside specifically in the coding region of the scrambling domain. Although it is currently unknown precisely how many ANO5 mutations affect scrambling, whether directly or indirectly through a reduction in protein expression, a loss of trafficking, or something else, it seems more than likely that the cell biology of ANO5 extends beyond its putative function as a PLSase. Our data suggest that one such role for ANO5 is as a facilitator of the translocation of key repair proteins following muscle wounding, though additional work is required to clarify whether this is through direct protein-protein interaction or some other mechanism.

## Methods

### Reagents

FM1-43 (T35356) was purchased from ThermoFisher. Cal520 AM (Cat # 21130) was purchased from AAT Bioquest. 1000X stocks of the dyes were prepared in distilled deionized water (FM1-43) or DMSO (Cal-520) and used within 7 days. Duramycin-Cy3 (Molecular Targeting Techonlogies D-1006) was prepared as a 500 μM (2000X) stock in 1% DMSO in deionized water.

### cDNAs

pEGFP-N1 (Clontech) was the backbone for many of the plasmids used in this study. This plasmid was modified by replacing EGFP with either NeonGreen (pmNeon), mEmerald (pmEmerald), tdTomato (pTomato), or mCherry (pmCherry). In some cases, proteins were tagged with a mutant GFP (A206K), which does not spontaneously homo-dimerize like WT GFP (81). GFP-A206K was generated through direct mutagenesis of the GFP sequence in the pEGFPN1 vector. A list of gene/tag combinations is provided in Table S1. We note no changes in the behaviors of the proteins studied when using different fluorescent protein tags. In addition to WT human ANO5, two mutants were used: an R58W mutant produced by point mutagenesis of WT ANO5, and chimeric “ANO5-1-5”, in which ANO5 amino acids 530-564 are replaced by amino acids 554-588 from ANO1. Production of ANO5-1-5 by replacing the scrambling domain of hANO5 with the homologous sequence from mANO1 has been described previously.

The cDNA construct for expression of Clover fluorescent protein conjugated to lactadherin C2 domain (LactC2-Clover) in the pET-28 bacterial vector was a gift from Dr. Leonid Chernomordik (NIH/NICHD). Purification of this probe has been described previously (72). The plasmid encoding LactC2-mCherry was made by excising Clover with XbaI and EcoRI and ligating in mCherry. This was purified identically to LactC2-Clover. Stock solutions (1-3 mg/ml) in 150 mM imidazole pH 7 plus 0.02% sodium azide were diluted to 2 μg/ml for PtdSer imaging.

### Mice

All procedures involving animals were approved by the Emory Institutional Care and Use Committee under protocols 201800130 (Hartzell) and 201700233 (Choo). *Ano5* KO mice have been described previously, and were bred and maintained as homozygous knockouts. Age-matched control C57BL/6 mice were purchased from Jackson Laboratories. Male mice between the ages of 3 and 6 months were used for all experiments. Female mice were excluded because Ano5-myopathies appear to disproportionately affect males (e.g. (82–84)).

### Electroporation of flexor digitorum brevis (FDB) muscle fibers

FDB fibers were transfected by *in vivo* electroporation. Mice were maintainted under isoflurane anesthesia throughout the procedure. Footpads were first injected with 20 μl 0.5 mg/ml hyaluronidase (H4272, Sigma-Aldrich). After two hours, < 20 μl of plasmid was injected into the footpad (total plasmid injected was typically 20 μg, but ranged from 10 to 50 μg). Electroporation was performed with a two-needle array electrode connected to an ECM830 *in vivo* electro-square porator (BTX Harvard Apparatus). 20 pulses (300 V/cm) lasting 20 ms were delivered at 1 second intervals. FDB muscles were isolated 7 days after electroporation to allow for muscle healing and plasmid expression.

### Isolation of muscle fibers

Mice were euthanized by isoflurane overdose followed by cervical dislocation. FDB muscles were removed, rinsed briefly in Hanks’ balanced salt solution, and transferred to digestion buffer consisting of 0.2% collagenase A (Roche 10103586001) in HEPES-buffered DMEM. Muscles were digested for 3 hours at 37°C, then transferred to DMEM containing 10% BSA. Individual fibers were dissociated via pipetting through a series of pipet tips with decreasing bore size. Following dissociation, fibers were allowed to settle for 15 minutes and were then resuspended in a vitamin-free modification of DMEM consisting of 1.8 mM CaCl_2_, 0.8 mM MgSO_4_, 5.3 mM KCl, 44 mM NaHCO_3_, 110 mM NaCl, 0.9 mM NaH_2_PO_4_, 1 mM sodium pyruvate, 5.6 mM D-Glucose and MEM amino acids solution (ThermoFisher 11130051, used at 2X final concentration) supplemented with 0.4 mM L-serine, 0.4 mM glycine, 4 mM L-glutamine, and 100 U/L penicillin/streptomycin (ThermoFisher 15140122). Fibers were seeded onto Matrigel (Corning 356234) coated glass-bottom dishes (MatTek, P35G-0-14-C) and allowed to adhere for at least 30 min prior to imaging. The skeletal muscle myosin II inhibitor N-benzyl-p-toluene sulphonamide (BTS, Tocris 1870) was added to fiber dishes at a concentration of 15 μM to reduce Ca^2+^ induced contraction after plasma membrane injury. All experiments were conducted within 24 hours of fiber isolation.

### Isolation and culture of human muscle progenitor cells (hMPCs)

All studies with human cells were reviewed and approved by the Emory Institutional Review Board (IRB00105208 and IRB00084168) and comply with all recommendations and requirements of the National Institutes of Health. Written informed consent was obtained from all subjects (except anonymous autopsy material). Human muscles were isolated by biopsy or autopsy. Isolated muscle chunks were minced by blades and incubated with 0.25% Trypsin for 20 minutes and filtered to isolate mononucleated cells including satellite cells. Cells were cultured on 1% porcine gelatin (Sigma G1890) -coated plates to expand and then sorted by surface markers (CD31^−^/CD45^−^/CD56^+^) using flow cytometry. Sorted hMPCs were grown on gelatin-coated dishes in Ham’s F10 medium supplemented with 10% fetal bovine serum (Sigma 12303C), 5% heat-inactivated calf-serum (Hyclone SH3008703), 0.5% chick embryo extract (US Biological C3999), 100 U/ml penicillin/streptomycin (Gibco 15140122) and 5 ng/ml basic fibroblast growth factor (Peprotech 100-18b). Cells were differentiated in low-glucose DMEM with 2% horse serum (Gibco 16050130) on gelatin coated plastic imaging dishes (ibidi). For injury experiments, subconfluent cells were differentiated for 14 days to produce elongated myotubes.

### Laser injury and imaging of membrane damage and repair

Culture dishes containing isolated muscle fibers were placed in a stage-top incubator (Tokai-Hit) heated to 37°C and gassed with 95% air/5% CO_2_. All images were taken on a Nikon A1RHD25 scanning confocal microscope using a 60x 1.4 NA oil objective. Digital zoom was set to 2.5 and a scanning resolution of 1024×1024 was used for all experiments, resulting in a pixel size of 0.08 μm. Pinhole size was set to 1 (myofiber injury) or 2 (hMPC injury) Airy units. A 2 μm x 2 μm (24.1pixel x 24.1pixel) region of interest was specified at the lateral edge of the fiber for laser ablation. Fibers were irradiated with a 405 nm laser set to 100% power (corresponding to ~0.9 - 1.1 mW) for 8 (myofibers) or 2 (hMPCs) seconds using the ND Stimulation toolbox in Nikon Elements software. Images were taken immediately prior to and after injury, then for every 4 s for a total of 2 minutes (30 frames), and finally every 15 s for a total of 5 minutes (21 frames). Times are reported as time from the start of scanning image #1 (pre-injury) t = 0.

#### Z Imaging

Z-stacks were acuqired in Nikon Elements software using the ND Acquistion menu. Highest and lowest focal planes of interest were identified by manually focusing through the depth of the injured region of the fiber. Images were taken every 0.5 μm with microscope and camera settings identical to those used to acquire timelapses after laser-induced injury.

#### Deconvolution

Images were deconvolved using the deconvolution module in Nikon Elements AR Analysis software. 2D deconvolution without subtraction was used for XY images, and the point spread function was determined automatically by the software. All quantification was performed on original images. Deconvolution is only for the purpose of presentation and clarity, and is noted in figure legends where applicable.

#### Ca^2+^ imaging

Cal-520 was mixed 1:1 with Pluronic F-127 and diluted to a final concentration of 10 μM in imaging medium with 25 mM HEPES. Fibers were loaded with Cal-520 in this solution for 30 minutes at RT, at which point the loading solution was replaced with fresh imaging medium. Fibers were then seeded onto MatTek dishes and allowed to adhere for 30 minutes prior to imaging.

#### FM1-43 repair assay

Imaging medium was replaced with imaging medium containing 2.5 μM FM1-43 at least 10 minutes prior to initiation of wounding experiments. For ANO5 rescue experiments, fibers expressing the desired fluorescent constructs were identified in the absence of FM1-43. This was done because FM1-43 has broad excitation/emission peaks in green and red wavelengths which would mask ANO5-expressing fibers if maintained in the bath at a working concentration. When ANO5-expressing fibers were identified, the X and Y positions of the microscope stage were marked using the ND Acquistion panel in Nikon Elements software. 100 μl of a 10X stock solution of FM1-43 in imaging medium was added dropwise to the dish (final buffer volume = 1 ml) with additional mixing by gentle pipetting. Fibers were left in the presence of FM1-43 at least 10 minutes prior to imaging.

### Quantification

All image analysis was performed on unaltered images using Fiji (ImageJ) software. Analaysis was not blinded with respect to experimental group.

#### Time courses

Elliptical ROIs were placed within the fiber boundaries and aligned to the center of the wound. Mean fluorescent intensity was measured within the ROI for each acquisition timepoint. Measurements were normalized to the mean fluorscence intensity from the first frame (F_0_) of the time series and plotted as a function of time in seconds. ROIs of the following sizes were used: 1000 μm^2^, mouse myofibers in the presence of FM1-43 or loaded with Cal-520; 50 μm^2^, hMPCs in the presence of FM1-43; 17 μm^2^, annexin electroporated mouse myofibers; 100 μm^2^, lipid probes.

#### Volumes

Reported volumes were calculated from z-stack images according to the formula

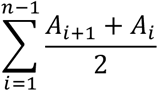

where A_*i*_ is the area of an object of interest, determined by manually encircling the object in ImageJ, from slice *i* of a stack of *n* images taken at 0.5 μm intervals.

#### Membrane blebs

Blebs were manually annotated from z-stack images in ImageJ using the multi-point tool. After scanning the entire stack, images were checked to ensure that blebs spanning more than one slice were not counted more than once.

#### Spatial localization of annexins in repair cap

ANXA1, ANXA5, and ANXA6 were co-electroporated into fibers with ANXA2. The final imaging frames from timelapse experiments were used for analysis (approximately 7.5 minutes after injury as described above). In ImageJ, the line tool was used to draw a line through the center of the ANXA2 cap. This line was set as an ROI so that its X and Y coordinates within the viewing frame in ImageJ were stored. The intensity profile in the ANXA2 channel was plotted for this line, then the intensity profile in the channel for the co-electroporated annexin along the same line was plotted. The absolute intensity data were transformed according to the formula

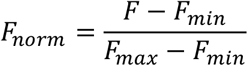

so that all values for each line intensity profile ranged between 0 and 1. F_norm_ values for both annexin proteins were plotted as a function of a variable termed here as “distance.” Units of distance are in μm and reflect points along the length of the line used to generate intensity profiles. For each replicate, distance = 0 was defined as the point where F_norm_ for ANXA2 [F_norm_(A2)] = 1. F_norm_ values for the co-electroporated annexin are plotted relative to the distance scale defined by ANXA2. F_norm_ values are shown for distances of −3 μm (towards myofiber center relative to F_norm_(A2) =1) to 3 μm (away from the myofiber center relative to F_norm_(A2) = 1). As an example, if the maximal F_norm_ value for the co-electroporated annexin protein was closer to the center of the fiber than F_norm_(A2), you would expect to see the curve of the co-electroporated annexin shifted to the left in a plot of F_norm_ vs. distance. The point (0,1) appears in every plot for ANXA2 because distance = 0 is defined by F_max_(A2), but because of variability from fiber to fiber, F_norm_(A[X]) may not equal 1.

#### Curve fitting

The time courses of protein accumulation were fit either to a single exponential equation or the sum of two exponentials using the fit functions in Origin 2019.

ANO5 and dysferlin were fit to an equation of the form

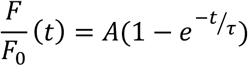

where A represents the amplitude (Y_plateau_ − Y_0_) and τ is the time constant.

Annexins were fit to an equation of the form

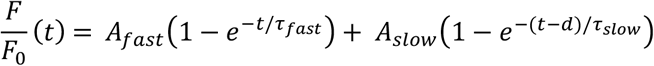

where A_fast_ and A_slow_ represent amplitude (Y_plateau_ − Y_0_) contributed by the fast and slow components, respectively, τ_fast_ and τ_slow_ are the time constants of the fast and slow components, and d is a time offset for the second exponential. R^2^ was >0.995 in all cases.

### Statistics

Statistical tests were performed in Prism 8 software (GraphPad). Data were analyzed via Student’s t-test or one-way analysis of variance (ANOVA) with Tukey’s post-test for multiple comparisons, as appropriate. Timelapse experiments and spatial localizations of repair cap annexins were analyzed through repeated measures two-way analysis of variance (ANOVA) with Geisser-Greenhouse correction and Sidak’s post-test correction for multiple comparisons. p-values < 0.05 were considered statistically significant.

## Acknowledgments

We thank Drs. Samya Chakravorty and Steven Goudy for assistance in procuring muscle biopsies. We thank Dr. Eunhye Kim for assistance with isolation of human muscle progenitor cells and Dr. Fang Wu for technical assistance. The cDNA construct for expression of Clover fluorescent protein conjugated to lactadherin C2 domain (LactC2-Clover) in the pET-28 bacterial vector was a gift from Dr. Leonid Chernomordik (National Institute of Child Health and Human Development). This research was supported in part by the Emory University Integrated Cellular Imaging Microscopy Core. Research reported in the publication was supported by grants from the National Institutes of Health as follows: the National Institute of Arthritis and Musculoskeletal Diseases under award numbers R01AR067786 (to HCH), R01AR071397 (to HC), and F32AR074249 (to SJF), National Eye Institute under award number R01EY114852 (to HCH), and National Institute of General Medical Sciences under award number R01GM132598 (to HCH). The content is solely the responsibility of the authors and does not necessarily represent the official views of the National Institutes of Health.

