## Supplementary material for "Defective Trafficking of Annexins to the Site of Injury in ANO5-Knockout Muscle Fibers": SI Appendix

**^1^**H. Criss Hartzell, Department of Cell Biology, 535 Whitehead Biomedical Research Building, 615 Michael St., Emory University School of Medicine, Atlanta, GA 30322 USA

^2^Hyojung J. Choo, Department of Cell Biology, 542 Whitehead Biomedical Research Building, 615 Michael St., Emory University School of Medicine, Atlanta, GA 30322 USA

**This PDF file includes:**

Figures S1 through S4

Tables S1 and S2

Legends for Supplemental Movies 1 to 7

Supplementary Methods

Supplementary References

**Other supplementary materials for this manuscript include the following:**

Supplemental Movies 1 to 7

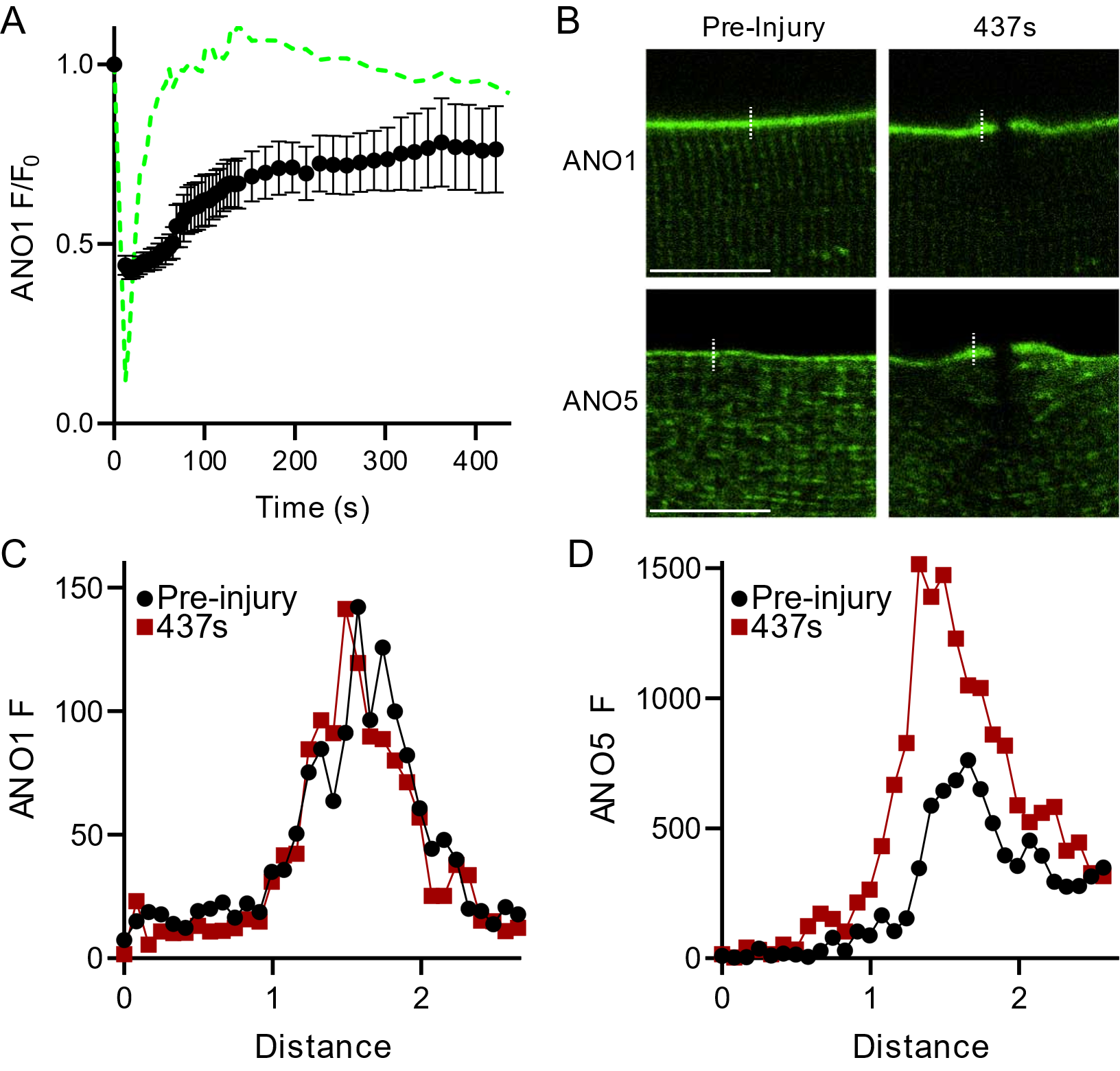

**Fig. S1.** Plasma membrane ANO1 does not traffic to wound sites despite localized fiber contraction. (A) Quantification of human ANO1 fluorescence at wound-adjacent sarcolemma following laser-induced injury of WT mouse myofibers. Fluorescence is normalized to pre-injury fluorescence values. Time course of human ANO5 accumulation in WT fibers (from Fig. 2) is shown as a green dotted line for comparison. ANO1 n = 10 fibers. Scale bar = 10 µm. (B) Images showing the contribution of localized myofiber contraction post-injury to the apparent accumulation of plasma membrane proteins at the wound site. (C) Line profile analysis of ANO1 fluorescent intensity before or after laser injury. Fluorescence values are measured from dotted lines as shown in top-row images of (B) mean fluorescence from n = 3 fibers. (D) Line profile analysis of ANO5 fluorescent intensity before or after laser injury. Fluorescence values are measured from dotted lines as shown in bottom-row images of (B) and are presented as mean fluorescence from n = 3 fibers.

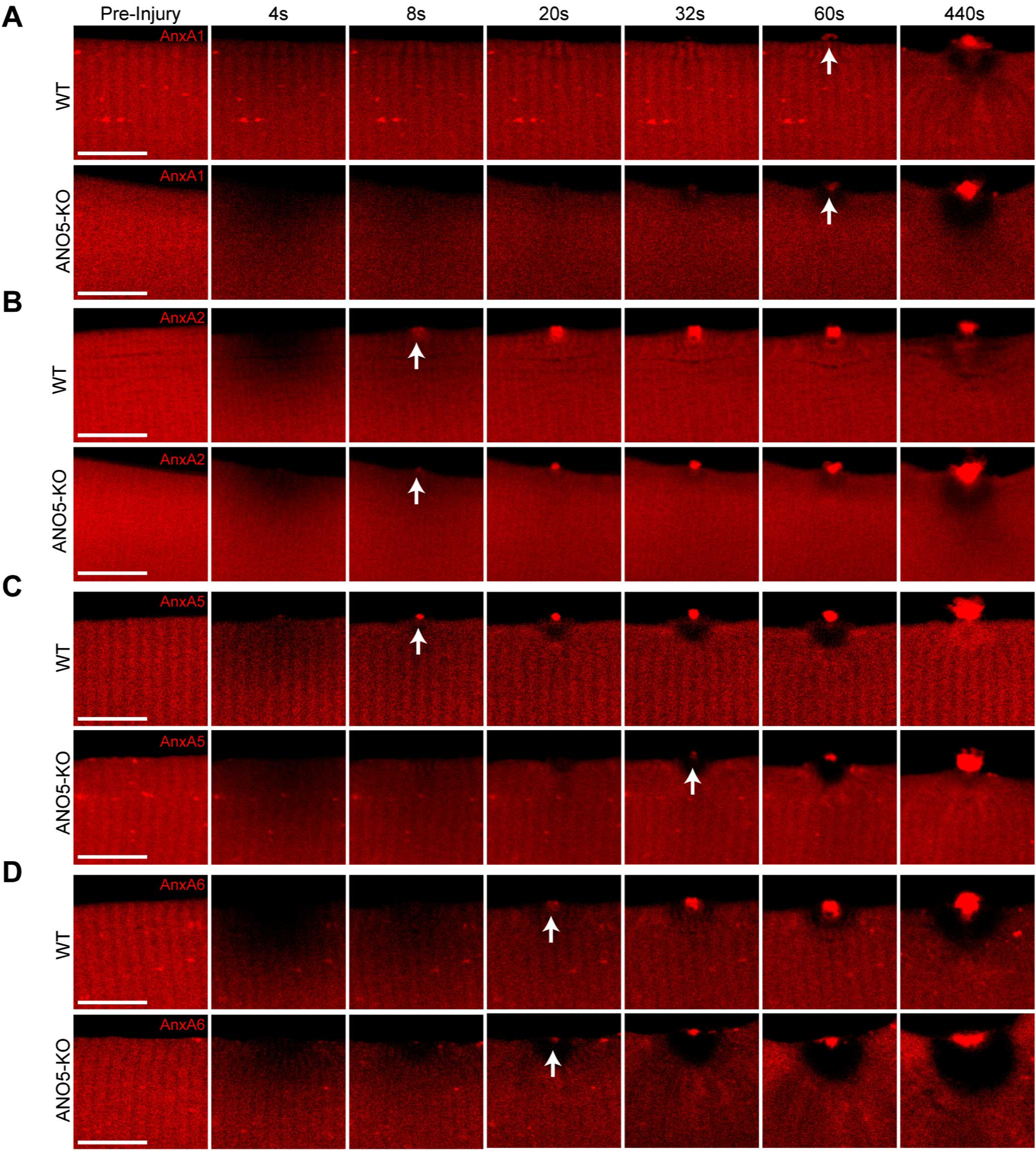

**Fig. S2.** Abnormal annexin trafficking and accumulation in ANO5-KO mouse muscle fibers. Representative images of (A) ANXA1, (B) ANXA2, (C) ANXA5 and (D) ANXA6 in WT or ANO5-KO fibers following laser-induced injury. Scale bar = 10µm. ANXA1: ANO5-KO n=13 fibers, WT n=14 fibers; ANXA2: ANO5-KO n=42 fibers, WT n=37 fibers; ANXA5: ANO5-KO n=36 fibers, WT n=29 fibers; ANXA6: ANO5-KO n=15 fibers, WT n=18 fibers.

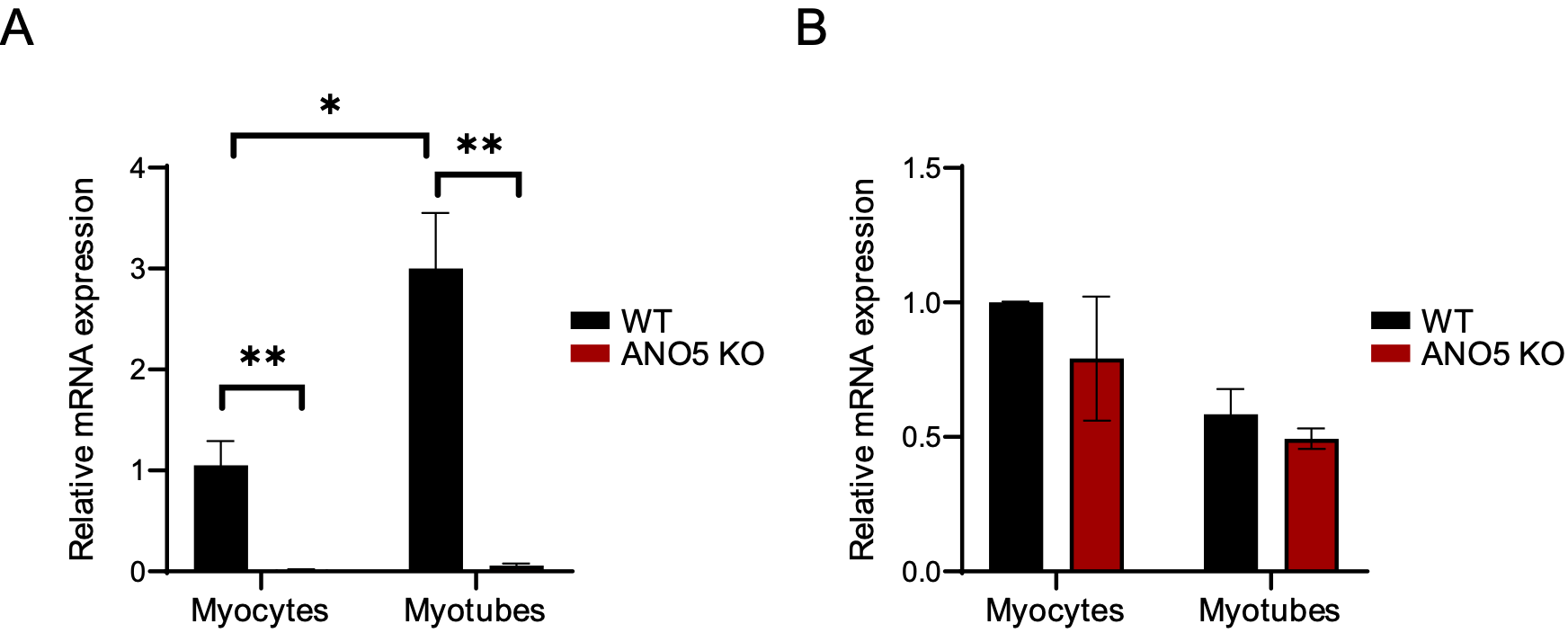

***Ano6***

***Ano5***

**Fig. S3.** *Ano5* but not *Ano6* mRNA is increased during myogenesis and *Ano6* expression is not changed in myocytes or myotubes from ANO5-KO mice. (A) mRNA expression of *Ano5* in WT or ANO5-KO mouse muscle precursor cells differentiated for 18 hours (myocytes, left) or 72 hours (myotubes, right). *, p < 0.05, ** p < 0.01. (B) mRNA expression of *Ano6* in in WT or ANO5-KO mouse muscle precursor cells differentiated for 18 hours (myocytes, left) or 72 hours (myotubes, right). WT n = 3, ANO5-KO n = 2.

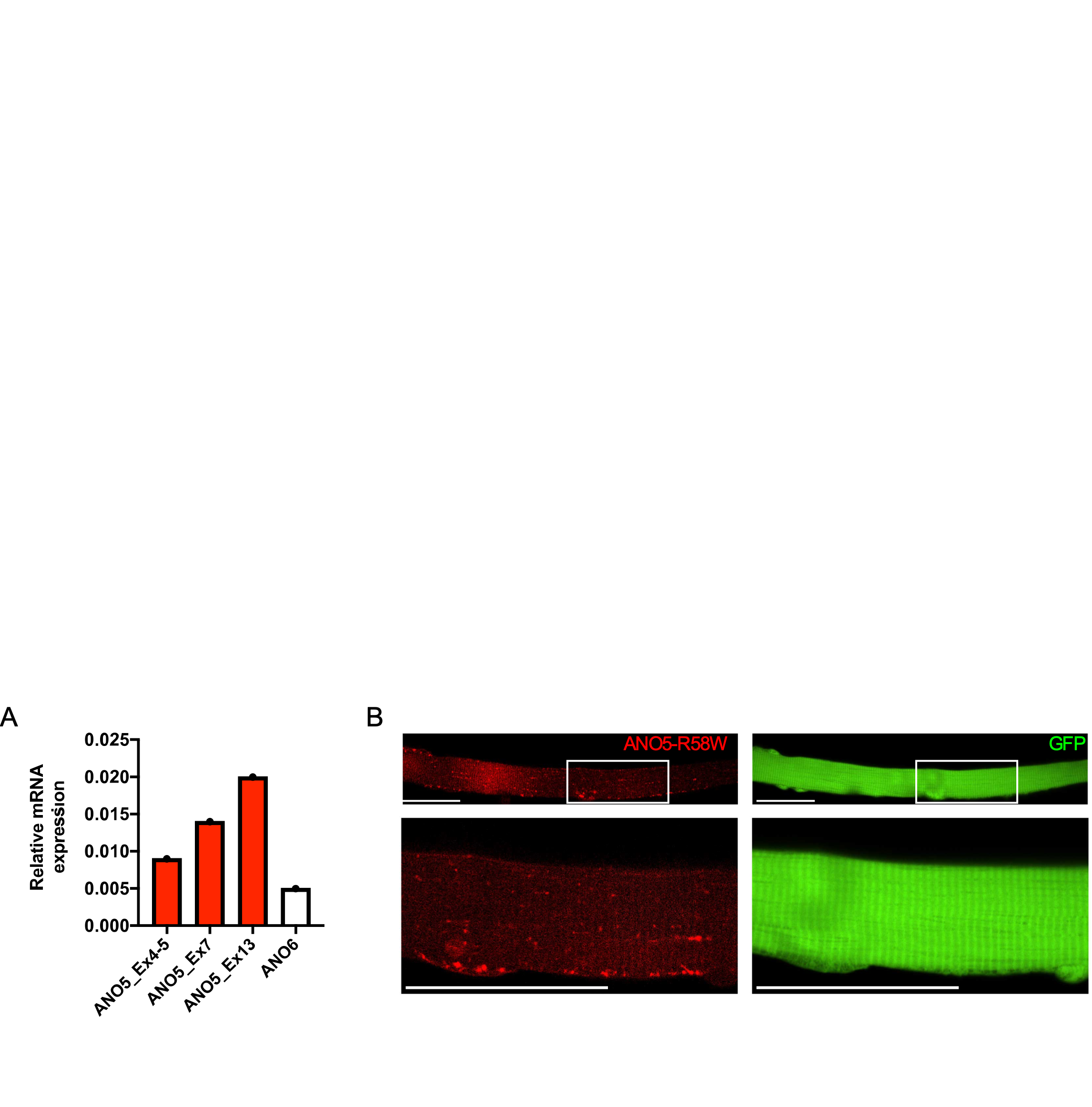

**Fig. S4.** Expression of ANO5 R58W mutant in human or mouse muscle. (A) qRT-PCR analysis from biopsied deltoid muscle of a human LGMD2L patient with primers directed at three regions. ANO5-R58W mutant does not produce degraded *ANO5* transcript. *ANO6* mRNA is shown as a comparison. (B) ANO5-R58W-mCherry is detectable when expressed in mouse muscle fibers. GFP alone is shown as a comparison.

| **Name** | **Uniprot/Gene Accession Number** | **Source** | **Lab of Origin** | **Fluorescent Protein Tags** |
| --- | --- | --- | --- | --- |
| ANO5  (codon optimized) | Q75V66 | DNA2.0 (Newark, CA) | Hartzell  (synthesized by DNA2.0) | mNeonGreen |
| ANO1 (a,c) splice variant | Q5XXA6 | Dr. Uhtaek Oh, Seoul National University | Oh | EGFP |
| Dysferlin  (transcript variant 2) | NM_001077694 | DNA 2.0 (Newark, CA) | Jain Foundation | GFP-A206K |
|  |  |  |  | tdTomato |
| AnnexinA1 TruORF | NM_000700 | Origene | I.M.A.G.E  Consortium | mCherry |
|  |  |  |  | mEmerald |
| AnnexinA2 | BC009564.1 | Dharmacon | I.M.A.G.E  Consortium | mCherry |
|  |  |  |  | mEmerald |
| AnnexinA5 TruORF | NM_001154 | Origene | I.M.A.G.E  Consortium | mCherry |
|  |  |  |  | mEmerald |
| AnnexinA6 TruORF | NM_001155 | Origene | I.M.A.G.E  Consortium | mCherry |
|  |  |  |  | mEmerald |
| R-GECO1.2 | Synthetic | Addgene | Robert Campbell |  |

Table S1. Genes expressed in mature mouse myofibers and the fluorescent tags attached to them.

|  | WT τ_fast_ | KO τ_fast_ | WT τ_slow_ | KO τ_slow_ | Fast A_WT_/A_KO_ | Slow A_WT_/A_KO_ |
| --- | --- | --- | --- | --- | --- | --- |
| ANXA1 | 15.7 ± 0.8 | 11.8 ± 0.4 | 268.7 ± 30.7 | 263.6 ± 15.8 | 1.21 | 1.36 |
| ANXA2 | 24.0 ± 0.6 | 24.6 ± 0.9 | 118.0 ± 46.3 | 146.4 ± 23.0 | 1.00 | 3.58 |
| ANXA5 | 12.8 ± 0.6 | 12.3 ± 1.3 | 142.8 ± 6.9 | 243.3 ± 21.5 | 1.44 | 0.95 |
| ANXA6 | 40.4 ± 3.1 | 28.8 ± 1.5 | 830.8 ± 144.4 | 1142.9 ± 187.9 | 1.96 | 1.71 |

**Table S2**. Fitted parameters of annexin time courses. Data are presented as mean ± standard error, calculated according to the Error Propagation Formula in Origin.

**Supplemental Movie 1.** Patch material is shed in WT myofibers.

**Supplemental Movie 2.** Prominent blebbing of repair patch in ANO5-KO myofibers.

**Supplemental Movie 3.** ANO5 accumulates at the wound shoulder throughout the Ca^2+^ transient.

**Supplemental Movie 4.** Repair cap ANXA2 is released in vesicles from WT myofibers.

**Supplemental Movie 5.** Rapid accumulation of PtdSer sensor LactC2-Clover and PtdEtn sensor duramycin-Cy3 at a wound site in Ca^2+^-free media.

**Supplemental Movie 6.** ANO5 but not ANO5-1-5 overexpression in ANO5-KO fibers reduces ANXA2 accumulation in the repair cap.

**Supplemental Movie 7.** ANO5 or ANO5-1-5 overexpression in ANO5-KO fibers improves ANXA6 trafficking during the wound response.

**Supplementary Methods**

### Isolation and differentiation of mouse primary myogenic progenitor cells (MPCs).

### Mononucleated cells were isolated from the hindlimb muscles as described previously (1). Isolated myogenic progenitor cells were cultured in Ham's F-10, 20% fetal bovine serum, 5 ng/ml basic fibroblast growth factor, 100 U/ml penicillin and 100 µg/ml streptomycin on collagen-coated plates for 3 or 5 days. Medium was changed every 2 days. For differentiation, MPCs were seeded at ECL coated tissue culture plates and cultured until 90% confluence. Then culture medium was changed to differentiation medium (2% horse serum in low glucose DMRM, 100 U/ml penicillin and 100 µg/ml streptomycin) and further cultured for 18 hours (myocytes) or 72 hours (myotubes).

### Real time-PCR analyses.

Total RNA was isolated from primary cultured myocytes or myotubes and deltoid muscle tissues biopsied from an ANO5 R58W patient using Trizol according to the manufacturer's protocol. The reverse transcriptase reaction was performed using 250 ng of total RNA/sample using random hexamers and M-MLV reverse transcriptase (Invitrogen). cDNA was amplified using the SYBR select master mix (Applied Biosystems) and 2.5 µM of each primer. All RNA samples were tested for DNA contamination by PCR. Amplified cDNA signals were detected and analyzed by StepOne software v2.2.2 (Applied Biosystems) using *Hprt* as internal control for mouse samples and *GAPDH* as an internal control for human samples. Fold change of gene expression was determined using the ∆∆Ct method (2). Three to four independent experiments were performed. Human primer sequences were:

*ANO5_Ex4-5*F: 5’- GCGGCGGCTTATGTTTCAAAA -3’

*ANO5_Ex4-5*R: 5’- CGCCTTTAACTCTGCGTCTTTC -3’);

*ANO5_Ex7*F: 5’- TATTCCCCGCCCTAAGCACA -3’

*ANO5_Ex7*R: 5’- AGAAGGTTGCCTGATCTTCGAT -3’

*ANO5_Ex13*F: 5’- TTTTGGAAACAACGACAAGCCA -3’

*ANO5_Ex13*R: 5’- ACCATACTGGTGACGACAAGAG -3’

*ANO6*F: 5’- AGCAAAGAAGTTTGTCATCC -3’

*ANO6*R: 5’- GAATGGACAAAGCCTATCAC -3’

*GAPDH*F: 5’- CTTTTGCGTCGCCAG -3’

*GAPDH*R: 5’- TTGATGGCAACAATATCCAC -3’

Mouse primer sequences were:

*Ano5*F: 5’- TCTTCCCACTGAGCACTTTC-3’

*Ano5*R: 5’-TGAGCATTCCTACACCAACC-3’

*Ano6*F: 5’- CTTATCAGGAAGTATTACGGC-3’

*Ano6*R: 5’-AGATATCCATAGAGGAAGCAG-3’

*Hprt*F: 5’-TCAGTCAACGGGGGACATAAA-3’

*Hprt*R: 5’-GGGGCTGTACTG CTTAACCAG-3’

**Ca^2+^-free lipid imaging.**

Fiber isolation, injury and imaging were similar to methods described elsewhere in the manuscript except for the imaging buffer used. Specifically, FDB muscle fibers were isolated from WT and seeded on uncoated glass-bottom dishes in extracellular solution containing (mM): 140 NaCl, 5 KCl, 1 MgCl_2_, 1 pyruvate, 25 glucose, 25 HEPES, 10 EGTA. Fibers were maintained in this Ca^2+^-free solution for at least 10 minutes prior to imaging, but for no more than 60 minutes total.
